# Temperature-Dependent Replication and Sensitivity to Innate Immunity of Human Coronavirus HKU1

**DOI:** 10.64898/2026.05.07.723210

**Authors:** Julian Buchrieser, Eva Thuillier, Jeanne Postal, Amélie Wileveau, Florence Guivel-Benhassine, Isabelle Staropoli, Nell Saunders, Delphine Planas, Jamie Sugrue, Vincent Bondet, Ignacio Fernández, François Bontems, Françoise Porrot, Chloé Petiot, Matthieu Prot, Martin Jungbauer-Groznica, Laurence Arowas, Vincent Michel, Catherine Blanc, Sophie Trouillet-Assant, Timothée Bruel, Félix A. Rey, Marie-Anne Rameix-Welti, Nicoletta Casartelli, Arnaud Fontanet, Michael White, Darragh Duffy, Etienne Simon-Lorière, Olivier Schwartz

**Author notes:** co-corresponding authors: JB and OS. Equal contribution (Eva Thuillier, Jeanne Postal, Amélie Wileveau).

## Abstract

The human coronavirus HKU1, causing common colds and occasionally severe illness, remains largely uncharacterized because it has not been successfully grown on immortalized cells. Here, we identified Caco2 cells overexpressing TMPRSS2, the HKU1 receptor, as being highly permissive to infection. HKU1 replicated efficiently, formed syncytia and released infectious progeny in these cells at 33°C, the temperature of the nasal cavity, but was attenuated at 37°C. Viral entry occurred similarly at both temperatures, but subsequent viral RNA synthesis was enhanced at 33°C. Released virions displayed higher stability at 33°C. In Caco2 and primary epithelial nasal cells, HKU1 was sensitive to interferons (IFN), but induction of IFN stimulated genes, such as IFN-Induced Transmembrane Proteins (IFITMs), was delayed at 33°C. Once expressed, IFITMs comparably inhibited HKU1 fusion at both temperatures. In contrast, SARS-CoV-2 robustly replicated at 37°C. Thus, cellular permissiveness, innate immunity and viral properties collectively explain why HKU1 replicates more efficiently at nasal temperature. Our results highlight temperature-sensitivity disparities between coronaviruses, likely associated to different pathogenic outcomes.

## Introduction

Human coronavirus (HCoV) HKU1 is one of four seasonal coronaviruses that circulate globally, causing common cold symptoms and occasional severe respiratory illness in vulnerable individuals^1^. First identified in 2005^2^, HKU1 has remained poorly characterized compared to other seasonal (OC43, NL63 and 229E) or epidemic coronaviruses (MERS-CoV, SARS-CoV-1 and SARS-CoV-2), due to significant challenges in developing culture systems and studying its replication dynamics^3–5^. HKU1 infections peak during winter seasons, with significant regional and age-based differences. The global seroprevalence of HKU1 is similar to that of other seasonal coronaviruses and may reach 75-95%^6–8^. Most individuals seroconvert during childhood^6^. Up to 20%–39% of children are infected every year with each HCoV^9^. There are three main HKU1 genotypes, HKU1 A, B and C. HKU1 A and B Spikes have 85% identity, with high conservation of the receptor-binding domain (RBD)^10–13^. HKU1 C is a HKU1 A/B recombinant, and its Spike shares 99% amino acid sequence identity with HKU1 B.

The molecular mechanisms governing HKU1 entry have been recently elucidated. The HKU1 Spike first binds through its N-terminal domain to 9-O-acetylated α2,8 linked disialoside on target cells^14–16^. The resulting conformational changes in the Spike triggers exposure of the RBD. We identified the transmembrane serine protease 2 (TMPRSS2) as a functional receptor for HKU1 Spike protein^13^. TMPRSS2 is expressed on airway epithelial cells, and primes coronavirus Spikes and other viral envelope glycoproteins, such as influenza hemagglutinin, enabling fusion^17,18^. Structural studies have elucidated the binding interface between HKU1 Spike RBD and TMPRSS2, showing that the interaction occurs with high affinity (in the 100 nM range)^19–23^. The HKU1 Spike interacts with TMPRSS2 at a well-defined groove, and key residues are conserved between HKU1 genotypes, indicating a conserved mode of receptor engagement. Binding of HKU1 Spike impairs TMPRSS2’s proteolytic activity, suggesting modulation during viral entry^19,20,24^.

So far, human airway epithelial cell (hAEC) cultures have been the only model used to study HKU1 replication. When differentiated at the air-liquid interface and maintained at 33°C, the physiological temperatures of the upper respiratory tract, this epithelium support HKU1 amplification^3–5,13^. HKU1 infects mainly ciliated cells, which express TMPRSS2 on their membrane and their ciliated protrusions^5,13,25^. HAEC cultures also support the replication of other HCoVs, with differential modulation by temperature. For instance, NL63 and 229E replication is attenuated at 37°C compared to 33°C. In contrast, the more pathogenic coronaviruses SARS-CoV-2 and MERS-CoV replicate at both temperatures, though SARS-CoV-2 replication may be enhanced at 33°C late in infection^26–28^. Temperature sensitivity of SARS-CoV-2 is also variant-dependent, suggesting that temperature may influence SARS-CoV-2 evolution^29,30^.

The differential viral replication at 33°C compared to 37°C is likely due to a combination of cellular and viral factors. At the cellular level, many parameters including transcription and protein expression, membrane fluidity, cycling, viability depend on the temperature and may directly or indirectly impact viral replication^31^. On the viral side, increasing temperature from 4°C to 33°C, 37°C or 39°C impacts SARS-CoV-2 Spike binding to its receptor and cell–cell fusion, modulating early steps of the viral cycle^29,32^. The stability, folding and enzymatic activity of viral polymerases and proteases may also be temperature-dependent^33–35^. Regarding innate immunity, the IFN-mediated antiviral and inflammatory responses are slower and weaker at 33°C than at 37°C^26–28^, leading to reduced restriction and higher viral titers in the upper airway. However, the impact of temperature on HKU1 replication has not yet been investigated.

Attempts to amplify and propagate HKU1 in immortalized cell lines (including RD, L929, HRT-18, LLC-MK2, Vero, or Calu3) have consistently failed^2,3^, indicating that specific requirements were absent in these cells. Here, we overexpressed TMPRSS2 in a panel of cell lines. We determined that human enteric Caco2 cells with elevated TMPRSS2 levels become remarkably permissive to both HKU1 A and HKU1 B clinical isolates. We took advantage of this novel system to amplify HKU1 in large quantities. We then studied viral replication and host responses at different temperatures in Caco2 cells and hAECs.

## Results

### Caco2 cells overexpressing TMPRSS2 are highly susceptible to HKU1

Previous efforts to cultivate HKU1 on cell lines have been unsuccessful^2,3,36^, but most of the tested cells lacked the viral receptor TMPRSS2. We thus examined whether overexpressing TMPRSS2 in a panel of cell lines would render them permissive to infection. To this aim, we isolated one HKU1 A and one HKU1 B strain from nasal swabs of infected individuals, by amplification on hAECs (Figures 1a and S1a). Sequencing confirmed the identity of the isolated viruses (Figures S1-3). We first tested Caco2 cells that we previously engineered to stably express high levels of TMPRSS2 (Caco2-TMP2+ cells)^13^ (Figure S4a). Upon infection at 33°C, Caco2-TMP2+ cells formed large Spike+ syncytia (Figure 1b) and allowed viral amplification.

**Figure 1.**
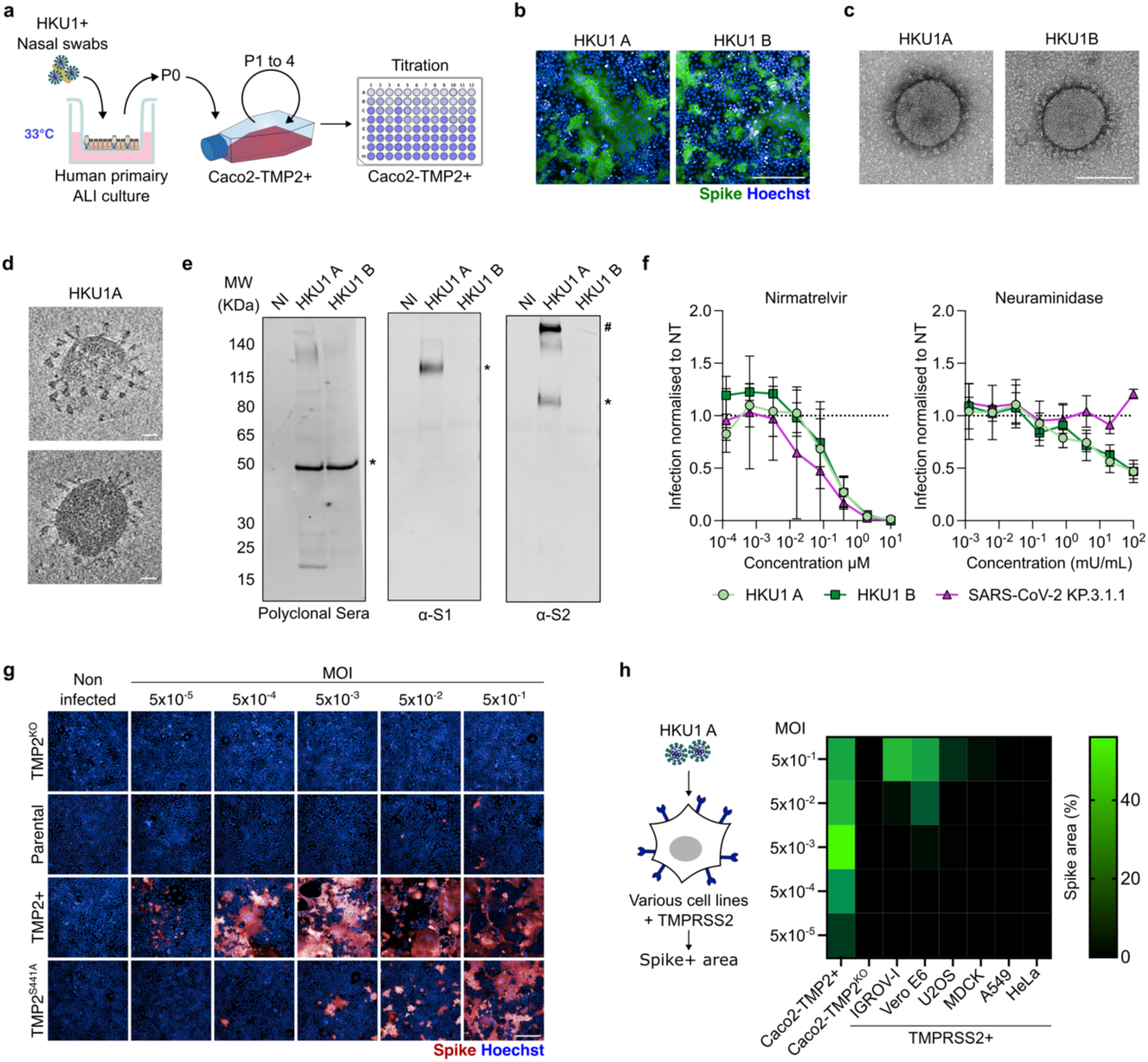
Caco2-TMP2+ cells are highly susceptible to HKU1. **a Amplification protocol of HKU1 in Caco2-TMP2+ cells.** Supernatant from primary hAEC culture was used to infect Caco2-TMP2+ cells at 33°C. Cultures were kept until cytopathic effect was detected. Supernatants were collected and the virus was further amplified on Caco2-TMP2+ cells. Viral stocks were sequenced and titrated on Caco2-TMP2+ cells. **b Caco2-TMP2+ cells form syncytia upon HKU1 infection.** Representative images of Caco2-TMP2+ cells infected with HKU1 A or B and stained for Spike 48h post-infection (pi). Green (Spike), Blue (Hoechst). Scale bar = 300 μm. **c Negative-stain transmission electron micrograph of HKU1 A and HKU1 B virions produced in Caco2-TMP2+ cells.** Virions display a characteristic morphology with Spike projections. Scale bar = 100 nm. **d Cryo-electron tomogram of HKU1 A particles produced in Caco2-TMP+ and purified by ultracentrifugation.** Surface Spike proteins are visible, projecting from the viral envelope. Scale bar = 25 nm. **e Western blot analysis of ultracentrifuged HKU1 A and B viral particles produced in Caco2-TMP+ cells.** Viral proteins were probed with three antibodies: (Left) Serum from an HKU-infected individual, the nucleocapsid protein (band labeled **\***, N, ∼50–55 kDa) generated the strongest signal. (Middle) Polyclonal antibodies against the S1 subunit of HKU1 A Spike (band labeled **\***, ∼120 kDa), (right) Polyclonal antibodies against the S2 subunit of HKU1 A Spike (band labeled **#**, corresponding to full-length S ∼220 kDa, and band labeled *, to the S2 fragments, ∼90 kDa). Representative blot of 3. **f Neuraminidase and Nirmatrelvir inhibit HKU1 A and B infection.** Caco2-TMP+ cells were pre-treated with indicated concentrations of Nirmatrelvir or Neuraminidase and infected with HKU1 A/B or SARS-CoV-2 KP.3.1.1 variant. Infection was measured after 48 h by Spike area quantification. Data are normalized to the non-treated condition, and are mean ± SD of 3-4 independent experiments. **g Levels of TMPRSS2 and permissivity of Caco2 cells to HKU1.** The indicated Caco2-derived cells were infected at various MOI with HKU1 A and analyzed four days pi. Representative images of infected Caco2 cells are shown. Red (Spike), blue (Hoechst). Scale bar = 300 μm. **h Relative susceptibility to HKU1 of various TMPRSS2 overexpressing cell lines.** Indicated cell lines were infected with a range of MOIs and the Spike area was quantified 4 days post infection. Heatmap of Spike area at indicated MOIs is shown. Data are mean of 2-3 independent experiments.

Viral stocks, after 3 passages on Caco2-TMP2+ cells, titrated up to 1x10^6^ TCID50/ml for HKU1 A and B (Figure S5a-b). Sequencing after 3 passages did not show signs of viral adaptation, except one fixed mutation for each isolate: HKU1 A nsp10 (L4377R) and HKU1 B nsp8 (K4065N), whereas K5969E in HKU1 A nsp14 was detected in about 45% of reads. Purified virions imaged by negative staining and electron microscopy (EM) exhibited the classical coronavirus aspect, size and distinctive crown-like Spikes protruding from the surface (Figure 1c). When imaged by Cryo-EM tomography, Spikes were visible, with shapes reminiscent of trimeric pre-fusion proteins (Figure 1d). Elongated post-fusion-like Spikes were also detected (Figure S6). Viral particles had a diameter of 107 nm ±13 and harbored 2 to 65 Spikes, with a mean of about 35 per particle (Table 1, Figure S6).

**Table 1.** Measurements of each cryo-electron tomogram of HKU1 A particle of Figure S5.

| **Virus n°** | **Spike**<br>**Number** | **Rotation**<br>**axis (nm)** | **Large axis**<br>**(diameter nm)** | **Eccentricity** | **Surface**<br>**(nm<sup>2</sup>)** | **Volume**<br>**(nm<sup>3</sup>)** |
| --- | --- | --- | --- | --- | --- | --- |
| Virus 1 | 65 | 60 | 110 | 0.84 | 27000 | 380000 |
| Virus 2 | 61 | 70 | 120 | 0.81 | 33000 | 530000 |
| Virus 3 | 57 | 60 | 120 | 0.87 | 31000 | 450000 |
| Virus 4 | 50 | 60 | 80 | 0.66 | 17000 | 200000 |
| Virus 5 | 49 | 70 | 120 | 0.81 | 33000 | 530000 |
| Virus 6 | 35 | 60 | 100 | 0.8 | 23000 | 310000 |
| Virus 7 | 34 | 60 | 110 | 0.84 | 27000 | 380000 |
| Virus 8 | 33 | 70 | 120 | 0.81 | 33000 | 530000 |
| Virus 9 | 28 | 40 | 100 | 0.92 | 20000 | 210000 |
| Virus 10 | 15 | 60 | 100 | 0.8 | 23000 | 310000 |
| Virus 11 | 12 | 60 | 100 | 0.8 | 23000 | 310000 |
| Virus 12 | 5 | 70 | 100 | 0.71 | 25000 | 370000 |
| Virus 13 | 5 | 60 | 100 | 0.8 | 23000 | 310000 |
| Virus 14 | 2 | 50 | 130 | 0.92 | 33000 | 440000 |
| Virus 15 | 2 | 60 | 100 | 0.8 | 23000 | 310000 |
| Mean | 30 | 61 | 107 | 0.81 | 26000 | 370000 |
| SD | 23 | 8 | 13 | 0.07 | 5200 | 110000 |

We then examined the virion protein content by western blotting. Samples were probed with a serum from an HKU1-infected individual and with rabbit anti-HKU1 A polyclonal antibodies targeting the Spike S1 and S2 subunits (Figure 1e). With the human serum, the nucleocapsid (N, ∼50–55 kDa) was detected with both HKU1 A and B particles, whereas Spike detection was minimal. The polyclonal antibodies recognized the cleaved forms of the HKU1 A Spike, S1 (∼120–130 kDa) and S2 (∼90–100 kDa), suggesting partial cleavage at the S1/S2 site. Uncleaved Spike (∼220 kDa) was detected with the anti-S2 but much less with the anti-S1 antibody. The HKU1 B Spike was not detected by these antibodies, as observed^13^.

We asked whether Nirmatrelvir (Paxlovid^TM^), a MPro inhibitor with antiviral activity against SARS-CoV-2, OC43 and 229E^37^, is active against HKU1. (Figure 1f). Nirmatrelvir potently inhibited HKU1 replication, with similar IC50 values as those observed for SARS-CoV-2 (KP.3.1.1 variant). We then analyzed the role of sialic acids during HKU1 infection. We treated Caco2-TMP2+ cells with neuraminidase, an enzyme that removes sialic acids. Neuraminidase inhibited HKU1 A and B, but not SARS-CoV-2 (Figure 1f), confirming that sialic acids are necessary for HKU1 entry.

We then asked whether the level and activity of TMPRSS2 affect susceptibility to HKU1 infection. We compared Caco2-TMP2+ cells to parental Caco2 cells (which express low levels of endogenous TMPRSS2), Caco2-TMP2^KO^ cells (that were knocked out for endogenous TMPRSS2), and Caco2-TMP2^S441A^ cells (lacking endogenous TMPRSS2 and expressing the catalytically inactive TMPRSS2-S441A mutant) (Figures 1g and S4a-b). As expected, Caco2-TMP2^KO^ were non permissive to HKU1. Parental Caco2 cells were only infected with high viral inoculum (MOI 0.5) while Caco2-TMP2^S441A^ were permissive, but to a lesser extent than Caco2-TMP2+. These results confirm, with replicative virus, our previous finding that TMPRSS2 enzymatic activity is dispensable for HKU1 entry^13^. Similar results were obtained with HKU1 B (Figure S4c). We then examined the antiviral activity of anti-TMPRSS2 nanobodies^13^. We tested the VHH A07 nanobody, which blocks the RBD interaction with TMPRSS2, and the VHH A01 nanobody which binds TMPRSS2 but does not prevent RBD binding^13^. VHH A07 blocked HKU1 infection in Caco2-TMP2+ and Caco2-TMP2^S441A^ cells while A01 increased it, likely by stabilizing the interaction between the two proteins^13^ (Figure S7). This confirmed that viral entry requires Spike-TMPRSS2 interaction in Caco2 cells.

Next, we tested the susceptibility to HKU1 of a panel of TMPRSS2-overexpressing cell lines (Figures 1h and S4d-f). Infected cells were detected at medium MOI (5x10^-3^) in IGROV-1-TMP2+ and VeroE6-TMP2+ cells. In U2OS-TMP2+ and MDCK-TMP2+ cells infection was only detected at a high MOI (0.5). Infection was almost undetectable even at high MOI in A549-TMP2+ and HeLa-TMP2+ cells. None of the parental cells were permissive to HKU1 infection. A549-TMP2+ and MDCK-TMP2+ cells were almost insensitive to infection, despite high levels of TMPRSS2 (Figures 1h and S4d).

Next, we assessed the level of sialic acids on a subset of these cell lines, using fluorescent *Sambucus nigra agglutinin* (SNA) Lectin (Figure S8a) that preferentially binds α-2,6- over α-2,3 linked sialic acids or recombinant mSiglec-E-Fc protein (Figure S8b) that preferentially binds α2,8-linked sialic acids^38^. Surprisingly, poorly permissive A549 cells express high levels of sialic acids levels which are reduced by neuraminidase, while Caco2 are poorly stained by the two tested lectins. Overall, variations in sialic acid levels do not account for the pronounced cell type–dependent differences in HKU1 replication. Thus, TMPRSS2 is necessary for HKU1 infection, with strong cell type-dependent variations in the sensitivity of target cell lines. Additional factors facilitate viral replication in Caco2 cells.

### Temperature sensitivity of HKU1 in cell lines

Following HKU1 isolation protocols on hAECs^3,36^, our infections were conducted at 33°C to reflect the upper airway temperature. We thus investigated the impact of temperature on HKU1 replication. We exposed Caco2-TMP2+ cells to HKU1 A and B at 33°C or 37°C. We followed overtime the appearance of infected cells and the release of infectious viral particles by TCID50 (Figure 2a) and release of viral RNA by qPCR (Figure S9b) in the extracellular medium. At 33°C, the number of infected cells increased rapidly over 72 h and large syncytia were visible with both HKU1 A and B. At 37°C, the infected cell area (Figure 2b), the released infectious virus (Figure 2c) and viral RNA (Figure S9b) were strongly reduced. Infection was strongly decreased with HKU1 A and almost undetectable with HKU1 B. In contrast, SARS-CoV-2 (KP.3.1.1 variant) replicated significantly better at 37°C than at 33°C (Figure S9a).

**Figure 2.**
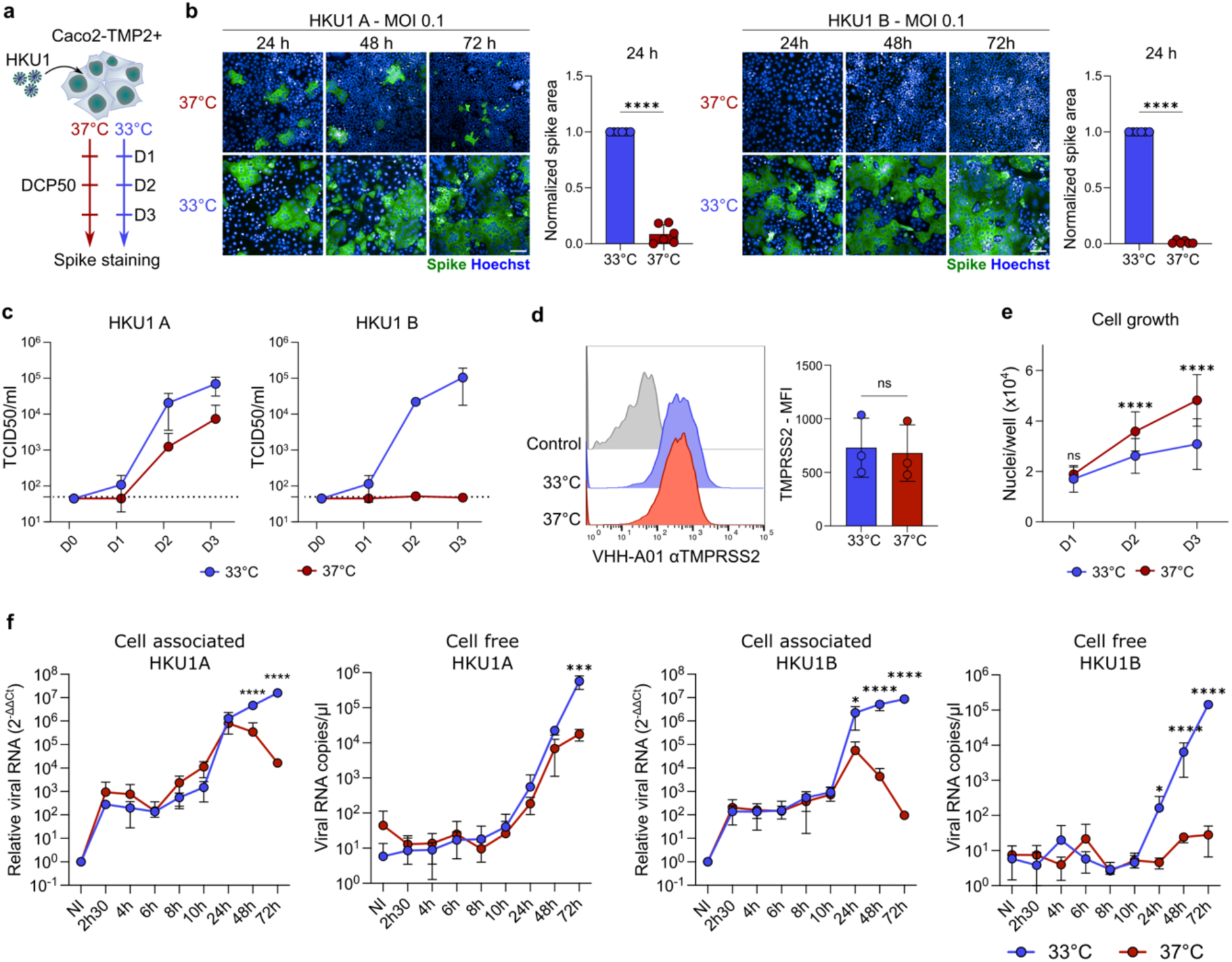
Temperature sensitivity of HKU1. A-D. HKU1 A and B replicate less efficiently at 37°C than at 33°C in Caco2-TMP2+ cells. **a** Caco2-TMP2+ cells were infected with HKU1 (MOI 0.1) at 33°C or 37°C for three days. Supernatants were collected for quantification of viral release and cells were fixed for Spike staining at the indicated days pi. **b** Representative images and Spike area quantification of Caco2-TMP2+ cells infected with HKU1 A (left) or HKU1 B (right) at 33°C or 37°C. Data are mean ± SD of 6 independent experiments. Scale bar = 100 μm. Statistical analysis: Unpaired t test with Welch’s correction. ****p <0.0001. **c Quantification of infectious viral release over time at 33°C and 37°C.** Data are presented as TCID50 and are mean ± SD of 2 independent experiments. **d Slower growth of Caco2-TMP2+ cells at 33°C.** Number of nuclei per well over time in non-infected cells at 33°C and 37°C. Data are mean ± SD of 4 independent experiments, with 9 analyzed wells per experiment. Statistical analysis: Two-way ANOVA with Šídák’s multiple comparisons test ****p<0.0001. **e TMPRSS2 surface levels at 33°C and 37°C.** Caco2-TMP2+ cells cultured for 48h at 33°C or 37°C were stained for TMPRSS2 using VHH-A01-Fc and analyzed by flow cytometry. (Left) Representative histograms of TMPRSS2 expression in cells cultured at 33°C (Blue) and 37°C (Red). Staining with secondary antibody alone is in Gray. (Right) Quantification of TMPRSS2 MFI. Data are mean ± SD of 3 independent experiments. Statistical analysis: Unpaired t-test, ns = non-significant. **f Cell-free and cell-associated viral RNA over time at 33°C and 37°C.** Caco2-TMP2+ cells were infected with HKU1 A or B for 2h30 and the virus inoculum was removed. Supernatant and cells were collected at the indicated timepoints pi and viral RNA was quantified by qPCR. Data are mean ± SD of 3 independent experiments. Statistical analysis: Two-way ANOVA with Šídák’s multiple comparisons test comparing 33°C to 37°C ***p<0.001, ****p<0.0001.

Next, we aimed to identify some of the cellular and viral factors that are modified at 33°C and could contribute to the observed differences in infectivity. Caco2-TMP2+ cell growth rate was significantly slower at 33°C than at 37°C, suggesting that this parameter may be involved (Figure 2d). In the infected conditions, no substantial cell death was observed at 37°C (Figure S9c-d). Flow cytometry staining showed no significant difference in surface TMPRSS2 levels in Caco2-TMP2+ cells cultured at 33°C or 37°C (Figure 2e). TMPRSS2 enzymatic activity, assessed with a fluorescent substrate, was unaffected in the 33-37°C range (Figure S10b). We also measured by flow cytometry the surface levels of HKU1 A/B or SARS-CoV-2 Spikes in transfected 293T cells and did not observe differences at the two temperatures (Figure S10c). We then examined the impact of temperature on Spike-mediated cell-cell fusion. We transiently expressed the HKU1 Spike and TMPRSS2 in 293T GFP-Split cells, which turn GFP+ upon fusion^39,40^. In line with results obtained with SARS-CoV-2 spikes^29^, fusion was significantly reduced at 33°C for HKU1 A/B and SARS-CoV-2 Spikes, when compared to 37°C (Figure S10d).

In contrast to SARS-CoV-2, HKU1 and OC43 encode for a hemagglutinin-esterase (HE) that has sialic acid receptor-destroying activity^14^ required for improved progeny virion release from infected cells. We thus investigated if the HE activity differed at 33 °C versus 37 °C. We transfected cells with HKU1 or OC43 HE for 24h and assessed the enzymatic activity of HE by incubating the cells in presence of a fluorescent substrate^41^ at 33°C or 37°C (Figure S11a). HE was highly expressed after transfection (Figure S11b) and an enzymatic activity was detected for both HKU1 and OC43 HE (Figure S11c-d). Temperature did not significantly impact the activity of the two HE (c-d).

We next measured the stability of cell-free HKU1 virions. We incubated viral particles at different temperatures for 1 to 4 h before measuring their infectivity. The virion infectivity declined significantly more rapidly at 37°C compared to 33°C (Figure S10e). After 4 h at 37°C virion infectivity was reduced by up to 60% for HKU1 A and 45% for HKU1 B (Figure S10f). Temperature may thus influence Spike stability, conformation or virion integrity.

Our next aim was to determine which steps of the HKU1 life cycle differed at 33°C and 37°C. We infected Caco2-TMP2+ cells and measured cell-associated and extracellular viral RNA over 72h (Figure 2f). With HKU1 A, cell-associated viral RNA started to increase at 8-10 h post infection (pi). Levels were similar at 33°C and 37°C, up to 24 h pi (Figure 2f). At later time-points, between 24 and 72 h pi, viral RNA exponentially grew at 33°C but dropped at 37°C. Cell-free viral RNA followed the same trend (Figure 2f). To determine background viral RNA levels in this system, we compared viral growth in Caco2-TMP2^KO^ and Caco2-TMP2+ cells at 33°C. Viral RNA copies above background were detected between 10 and 24 h pi for cell-associated and cell-free RNA (Figure S9g). With HKU1 B, the temperature dependence was more pronounced, with a significant difference between 33°C and 37°C in both cell-associated and free viral RNAs observed as early as 24 h. Thus, entry and initial viral RNA replication are minimally affected by temperature. The temperature dependence appears to be linked to either a late-stage replication block or to variations in cellular responses which may affect secondary infection of bystander cells.

We next infected IGROV-1-TMP2+, VeroE6-TMP2+ and A549-TMP2+ cells with HKU1 A or HKU1 B, at 33°C or 37°C (Figure S9e-f). The number of HKU1 A and B infected cells was also drastically reduced at 37°C, indicating that the low temperature dependance of HKU1 is not limited to Caco2 cells.

Overall, our results indicate that temperature influences several cellular and viral parameters that may facilitate HKU1 replication at 33°C in cell lines.

### A delayed IFN response enhances HKU1 replication at 33°C in primary hAECs

Temperature sensitivity of other human common cold coronaviruses (NL63, OC43, 229E) has been linked to a delayed and dampened cellular IFN response at 33°C in primary cells^28^. The underlying mechanisms, particularly the IFN stimulated genes (ISGs) involved, are poorly characterized. We thus asked whether HKU1 replication is dependent on the temperature in primary cells and examined the role of the IFN response in this model. We pre-incubated nasal hAECs at 33°C or 37°C with or without the JAK-STAT pathway inhibitor Ruxolitinib (10 μM) and infected them with HKU1 A. Virus release was measured by RT-qPCR and cytokine production was quantified using a Luminex multiplex assay over 4 days, after which the proportion of infected hAECs was determined by immunofluorescence (Figure 3a). At 33°C, 4 days pi, approximately 15% of cell area was infected (Spike positive), irrespective of Ruxolitinib treatment (Figure 3b-c). In contrast, at 37°C, only ∼1% of the cell area was infected, and Ruxolitinib increased infection (∼8% of Spike positive area) (Figure 3c). Accordingly, viral RNA release at different time points was attenuated at 37°C compared to 33°C (days 2-4 pi, Figure 3d). This decrease was no longer observed with Ruxolitinib.

**Figure 3.**
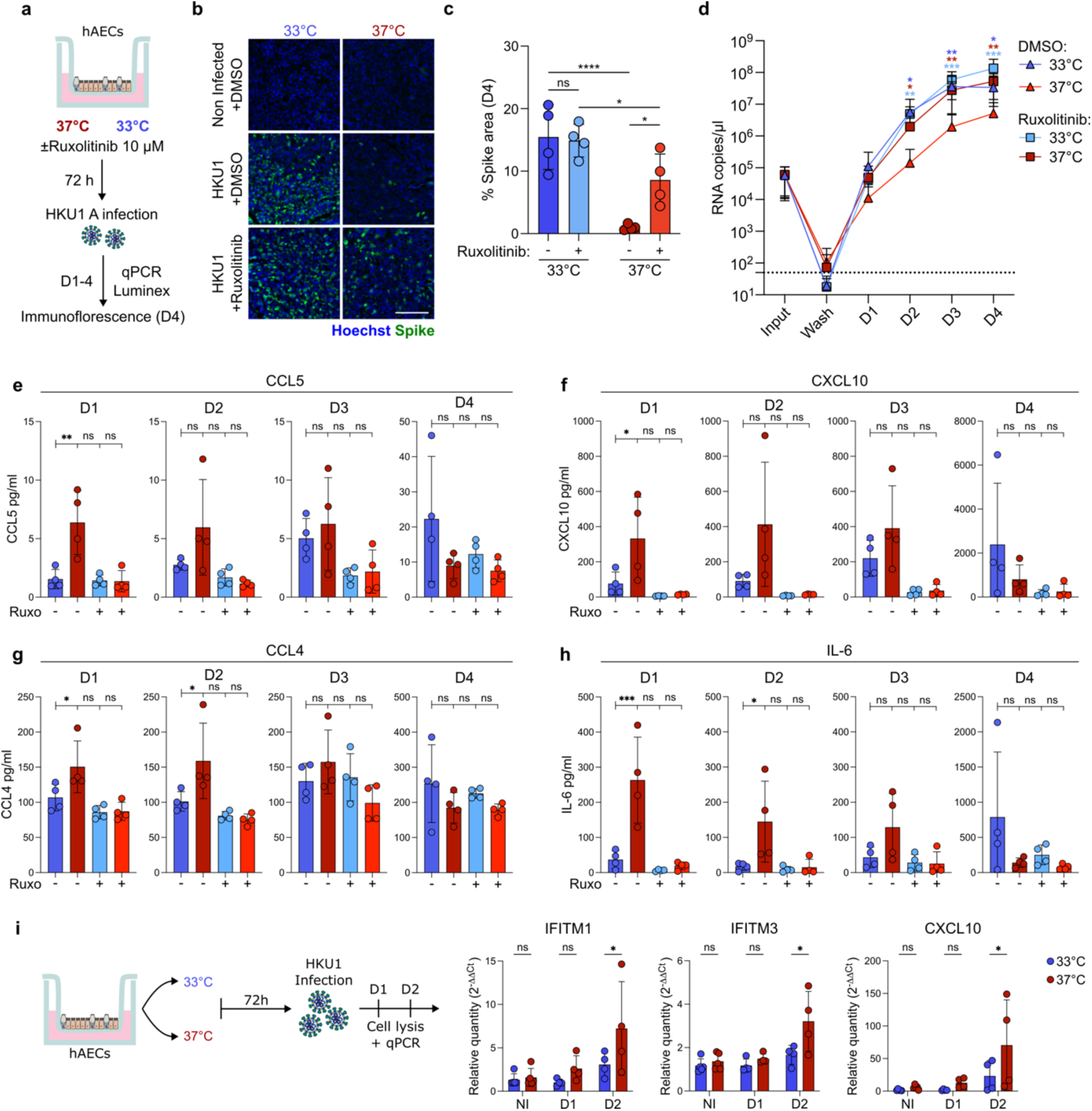
HKU1 replicates efficiently at 33°C due to a delayed ISG response in primary hAECs. **a** Experimental design: primary hAECs were treated in the basal chamber with DMSO or Ruxolitinib (10 μM) and incubated at 33°C or 37°C. After 72 h, cells were infected with HKU1 A and infection was followed over 4 days. **b Representative images of hAECs at day 4 pi.** Spike (Green), Hoechst (Blue). Scale bar = 100 µm. **c Ruxolitinib partially rescues HKU1 replication at 37°C.** The infection was quantified using ImageJ. Approximatively ¼ of a transwell insert was imaged per condition. Data are mean ± SD of 4 independent experiments. Statistical analysis: Two-Way ANOVA comparing cell means with others in its row and its column * p<0.05, ****p<0.0001. **d Quantification of HKU1 viral release.** Viral RNA copies in the supernatant of hAECs were quantified by RT-qPCR. Data are mean ± SD of 4 independent experiments. Statistical analysis: Two-Way ANOVA on the log_10_ transformed data with Tukey’s multiple comparisons test, comparing the 37°C DMSO condition to: 33°C DMSO (light blue stars), 33°C Ruxolitinib (Dark blue stars) or 37°C Ruxolitinib (Dark red stars). * p<0.05, **p<0.01, ***p<0.001. **e-h HKU1 infection induces a rapid ISG response at 37°C.** Cytokines in the apical compartment of hAECs was quantified over time by Luminex multiplex assay. CCL5 (E), CXCL10 (F), CCL4 (G) and IL-6 (H) are plotted. Data are mean ± SD of 4 independent experiments. Statistical analysis: One-Way ANOVA ns = non-significant, * p<0.05, **p<0.01. **i Antiviral ISGs upregulated upon infection at 37°C.** Primary hAECs were infected with HKU1 at 33°C or 37°C, cells were lysed at day 1 or 2 and ISG induction was quantified by qPCR. Relative quantities of IFITM1, IFITM3 and CXCL10 mRNA normalized to GAPDH endogenous control are plotted. Data are mean ± SD of 4 independent experiments. Statistical analysis: Paired Two-Way RM ANOVA * p<0.05, **p<0.01.

We examined the impact of temperature on the surface expression of TMPRSS2 and other coronavirus receptors. HAECs were cultured for 3 days at 33°C or 37°C, dissociated, and analyzed by flow cytometry (Figure S12a). Temperature did neither change the proportion of ciliated cells (β-tubulin positive) (Figure S12b) nor surface levels of TMPRSS2, ACE2, APN, and DPP4 (Figure S12c-h). We further studied by immunofluorescence the global localization of TMPRSS2 in hAECs. In agreement with previous results^13,25^, TMPRSS2 accumulated in ciliated protrusions, without major changes at 33°C or 37°C, with and without Ruxolitinib (Figure S13a).

To assess whether HKU1 A induces an ISG response in hAECs, we stained the cells for MX1, a known ISG^42^. MX1 was significantly upregulated at day 4 pi (Figure S13b). Ruxolitinib blocked MX1 upregulation. We further analyzed cytokine release by Luminex at different time points (Figure 3e-h). Three interferon-stimulated cytokines^42–44^, CCL5 (RANTES), CXCL10 (IP-10) and CCL4 (MIP-1β), were significantly more produced at 37°C compared to 33°C, at 24 h pi. Higher levels of IL-6, which signals through the JAK-STAT pathway^45^, were released at 37°C than at 33°C, at 24 and 48 h pi. At 33°C, CCL4, CCL5, IL-6 and CXCL10 release increased more gradually, surpassing at days 3 or 4 pi the levels detected at 37°C. Ruxolitinib dampened release of the four cytokines (Figure 3e-h). Other tested cytokines showed no major differences between conditions (Figure S14).

We also examined the induction of ISGs within HKU1-infected hAECs. We measured by RT-qPCR the RNA levels of CXCL10, and of IFITM1 and IFITM3, two proteins which restrict the replication of many viral species^46^. At 48 h pi, these ISGs were increased at 37°C compared to 33°C (Figure 3I), whereas viral RNA levels were higher at 33°C (Figure S13c).

Altogether, our results show that HKU1 induces an ISG response in primary cells, which is delayed at 33°C despite a stronger infection. Inhibiting the ISG response at 37°C with Ruxolitinib significantly enhances HKU1 replication.

### IFN response in Caco2 cells at 33°C and 37°C

We next examined the IFN response in Caco2 cells. The addition of IFN-β1b or IFN-λ induced IFITM1 and IFITM2 proteins (Figure S15a). IFN-β1b and IFN-λ also potently inhibited the replication of HKU1 A/B and SARS-CoV-2 (D614G and BA.1 variants). IFN-β1b upregulated various ISGs (STAT1, OAS1, LY6E, IFITM1/3 and MX1) significantly faster at 37°C than at 33°C (Figure S15b). Strikingly, in contrast to primary hAECs, Ruxolitinib did not rescue HKU1 A nor B infection at 37°C (Figure S16a-b) and had a minor effect on SARS-CoV-2 (KP.3.1.1 variant). Ruxolitinib was however active in Caco2-TMP2+ cell. It blocked the up-regulation of IFITM1 and IFITM2/3 induced by exogenous IFN-β1b (Figure S16d). We thus asked whether HKU1 infection upregulated ISGs in Caco2-TMP2+ cells. No induction of IFITMs was detected in infected cells at 33°C or 37°C (Figure S17a).

Next, we tested whether HKU1 can suppress IFN responses in Caco2-TMP2+ cells. Caco2-TMP2+ cells were infected with HKU1 A or B for 24h at 33°C and subsequently treated with IFN-β1b (1000 U/ml) for 24h at 33°C or 37°C (Figure S17b). Cells were stained for S and IFITM1 or IFITM2/3 and mean intensity of IFITM staining was quantified in the S positive area and bystander S negative cells (Figure S17c-f). IFITM1 and IFITM2/3 were significantly upregulated upon IFN treatment. We did not detect any significant difference in IFITM level in infected and bystander cells at 33°C or 37°C for both HKU1 strains. Therefore, in Caco2-TMP2+ cells, HKU1 does not markedly suppress the ISG response induced by IFN treatment.

In sum, IFN-β1b or IFN-λ upregulate ISGs, including IFITMs, in Caco2-TMP2+ cells. This induction is delayed at 33°C. This ISG response blocks HKU1 and SARS-CoV-2 infection. However, in Caco2-TMP2+ HKU1 infection does not trigger an IFN dependent antiviral response nor does HKU1 suppress ISG induction. Ruxolitinib does not rescue the attenuation of HKU1 replication at 37°C. Therefore, in Caco2-TMP2+ cells, IFN-independent mechanisms are mainly involved in the temperature dependence of HKU1.

### Exploring IFITM restriction of HKU1

Since IFITMs were upregulated upon HKU1 infection in hAECs (Figure 3i), potentially making bystander cells resistant to secondary infections, we asked whether these proteins are active against HKU1. We included LY6E, which also blocks coronavirus fusion^47,48^ and was upregulated by IFN in Caco2-TMP2+ cells (Figure S15). We first focused on the viral entry step by using lentiviral pseudotypes bearing the HKU1 Spike^13^. 293T cells were co-transfected with plasmids expressing TMPRSS2 and IFITM1-3 or LY6E and infected with HKU1 A and B pseudotypes encoding the luciferase reporter gene (Figure 4a). The luciferase signal was reduced at 33°C compared to 37°C (Figure S18c), likely because lentiviral replication or luciferase activity is optimal at higher temperatures. We thus normalized at each temperature the infections to the condition without ISG. IFITM1 and LY6E significantly inhibited HKU1 A and B pseudovirus infection at 33°C and 37°C (Figure 4a) without affecting TMPRSS2 surface levels (Figure S18d).

**Figure 4.**
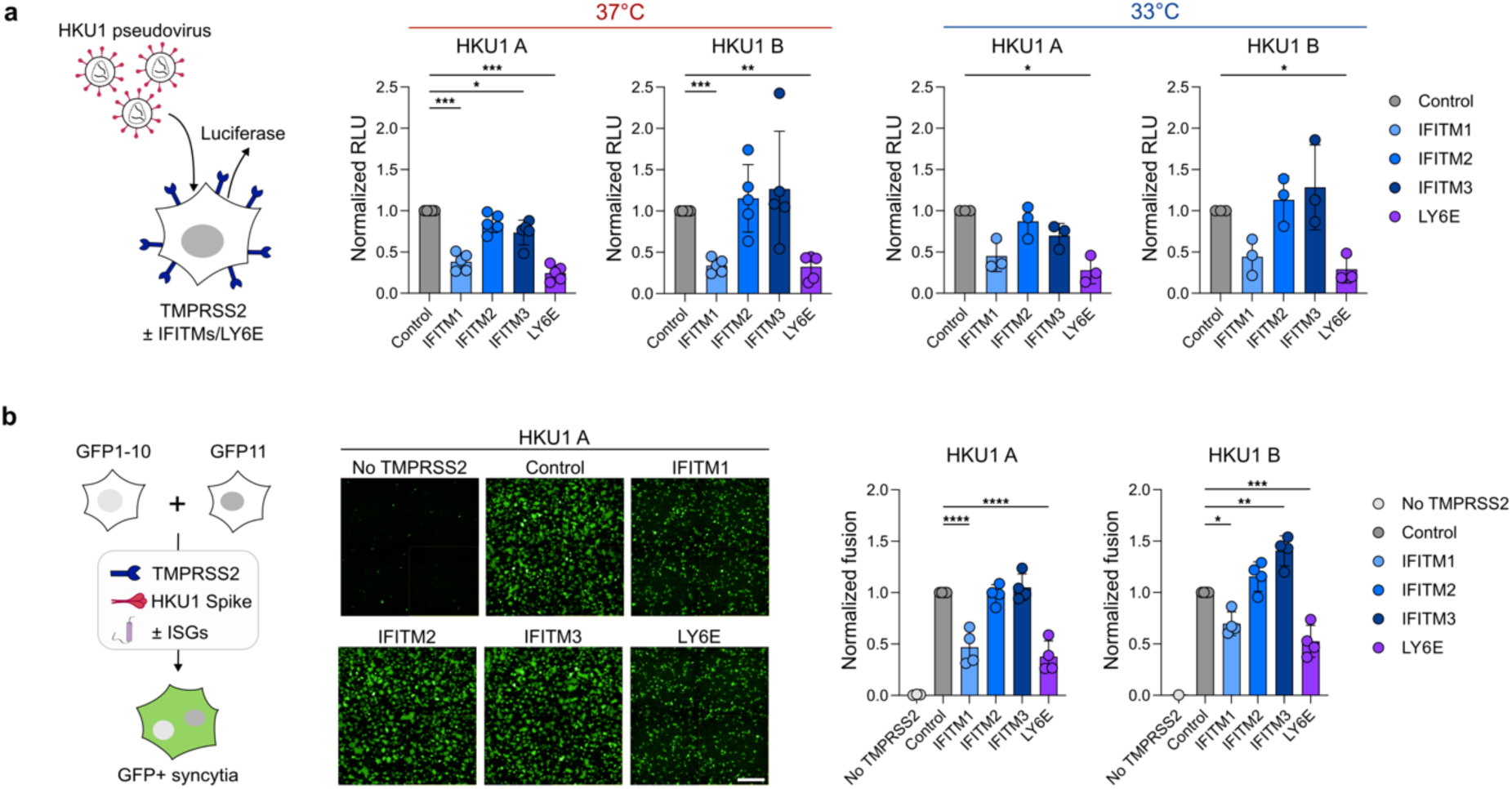
a IFITM1 and Ly6E inhibit HKU1 fusion and entry. 293T cells were transfected with TMPRSS2 and a control (Empty plasmid), IFITM1-3 or LY6E expression plasmids and maintained at 37°C. 24 h post transfection, cells were infected with luciferase encoding lentiviral HKU1A or B pseudotypes and placed at 33°C or 37°C for an additional 24 h. Normalized luciferase activity (RLU) at 37°C (left) and 33°C (right) are plotted. Data are mean ± SD of 5 (37°C) or 3 (33°C) independent experiments. Statistical analysis: One Way ANOVA with Tukey’s multiple comparisons * p<0.05, **p<0.01, ***p<0.001. **b Impact of IFITMs and Ly6E on HKU1 Spike mediated cell-cell fusion.** 293T-GFP1-10 and -GFP11 cells (1:1 ratio) were co-transfected with HKU1 A or B Spike, along with TMPRSS2, IFITMs, LY6E or control plasmids. Representative images are shown (middle). Scale bar = 500 µm. Cell fusion was quantified by measuring the GFP+ area by high-content imaging after 18 h (right). Data are mean ± SD of 4 independent experiments. Statistical analysis: One Way ANOVA with Tukey’s multiple comparisons * p<0.05, **p<0.01, ***p<0.001, ****p<0.0001.

We then assessed the effect of IFITMs and LY6E on HKU1-Spike mediated cell-cell fusion, using the 293T GFP-Split reporter system (Figure 4b). Fusion mediated by HKU A and B Spikes was inhibited by IFITM1 and LY6E, but not by IFITM2/3. Co-transfection of IFITMs slightly reduced HKU1 A/B Spike surface expression (Figure S18e). SARS-CoV-2 (D614G variant)-mediated Spike fusion was also inhibited by IFITM1, and to a lesser extent by IFITM2/3 and LY6E. This inhibition was reverted by TMPRSS2 (Figure S18b), as we previously described^40^. LY6E inhibited SARS-CoV-2 fusion irrespectively of TMPRSS2.

Thus, IFITM1 and LY6E inhibit HKU1 Spike-mediated cell-cell fusion and HKU1 pseudovirus entry. These ISGs are rapidly induced at 37°C in hAECs and may therefore contribute to HKU1 inhibition at this temperature.

### HKU1 A and B seroneutralization

Most of the available sero-epidemiology data regarding HKU1 have been obtained with viral proteins and peptides as targets, or sometimes with pseudovirus-based assays^9,49,50^. Seasonal coronavirus protective immunity is short lasting^51^, but little is known about the neutralization activity present in the sera of different categories of individuals. To assess this activity, we designed a neutralization assay using infectious HKU1 A or B virus and Caco2-TMP2+ cells. We selected sera from 100 individuals from the SeroPed cross-sectional study, which was implemented to evaluate immunity to human coronaviruses in individuals attending French hospitals^9^. These sera have been previously tested with a 9-plex assay to measure antibodies to 5 SARS-CoV-2 antigens and the Spike ectodomains of the 4 HCoVs (NL63, 229E, OC43, and HKU1 B). We picked 100 samples, ensuring representation of the entire spectrum of antibody responses and patient ages (Table 2). As expected, for both HKU1 A and B, neutralization titers, as well as the proportion of individuals positive for sero-neutralization, increased with age (Figure 5a). The correlation between neutralization of HKU1 A and B was weak (Figure 5b), suggesting partial cross-reactivity between the strains. We also compared the neutralizing antibody (NAb) titers to the levels of Spike-binding antibodies measured by Luminex^9^ (Figure 5c). The sera with high NAb levels were generally positive by Luminex. However, many Luminex-positive sera did not display NAb activity. This suggests that non-neutralizing antibodies may accumulate in infected individuals, or that the sensitivity of the neutralization assays is different from that of the Luminex assay.

**Figure 5.**
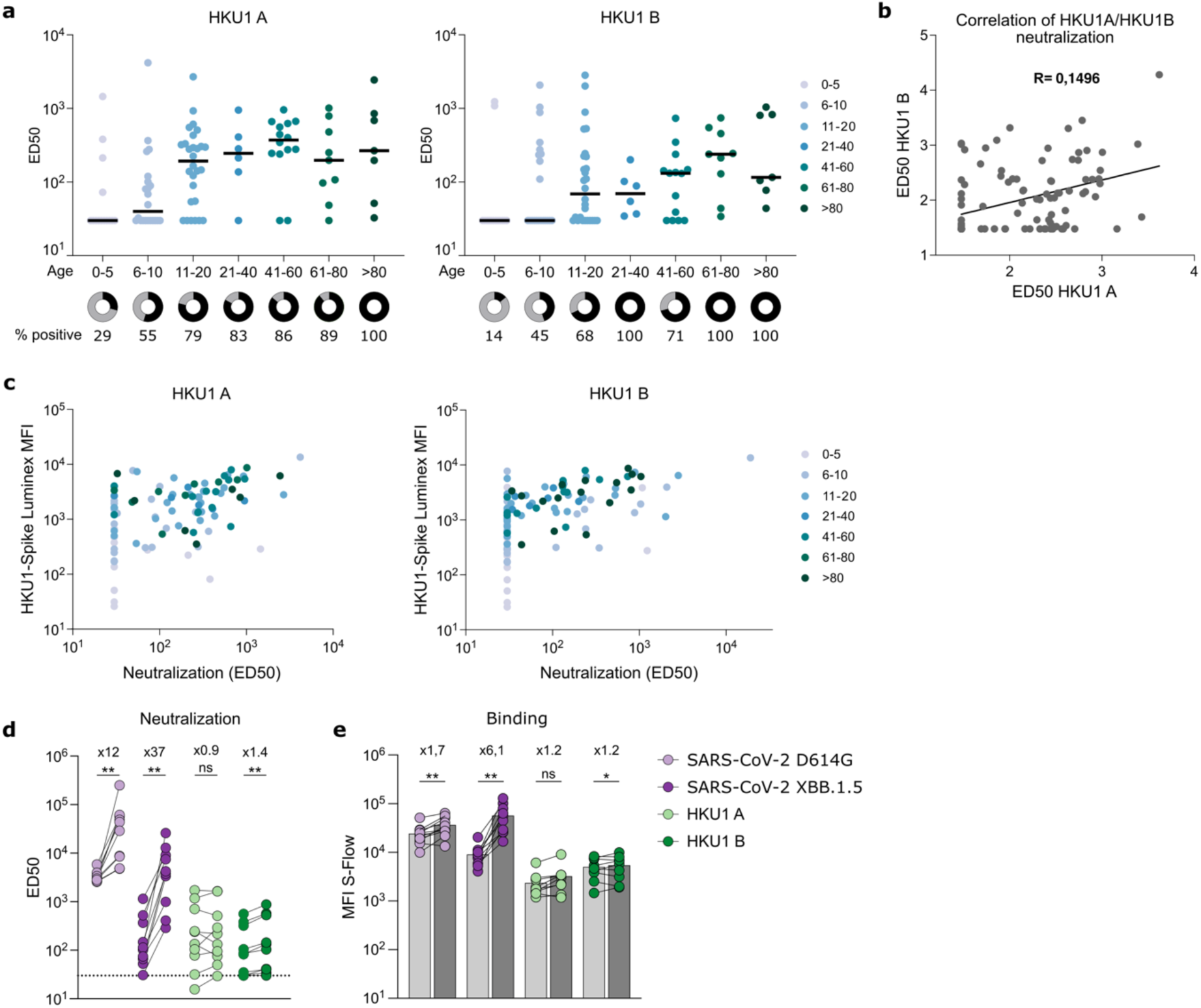
a Neutralization titers of sera from individuals of different age groups against HKU1 A (left) or HKU1 B (right). Neutralization titers are presented as ED50. The percentage of individuals positive for neutralization (above threshold of 30, corresponding to the first testeddilution) are represented as pie charts under their respective age group. Sera (*n* = 100). **b Limited correlation between HKU1 A and B neutralization.** Results obtained for HKU1 A were correlated with those of HKU1 B. Each dot represents one serum. Correlation coefficient (R) is shown. **c Correlation between HKU1-Spike Luminex results and HKU1 neutralization from individuals of different age groups.** Neutralization titers, presented as ED50 are plotted against the previously published^9^ MFI of HKU1-Spike Luminex results for HKU1 A (left) and HKU1 B (right). **d SARS-CoV-2 XBB.1.5 vaccination does not strongly boost HKU1 cross-neutralization.** Neutralizing activity of sera (n= 10) from individuals before (left) and one month after (right) XBB.1.5 booster dose. Neutralization titers against the indicated SARS-CoV-2 variant or HKU1 strain are expressed as ED50. Statistical analysis: Wilcoxon matched-pairs signed rank test, ns = non-significant, **p<0.01. **e Binding of sera from individuals before and one month after XBB.1.5 booster dose.** Data are plotted as MFI. Sera were tested at a 1:100 dilution. Data are the mean of 2 independent experiments. Statistical analysis: Wilcoxon matched-pairs signed rank test, ns = non-significant, *p<0.05, **p<0.01.

**Table 2.**
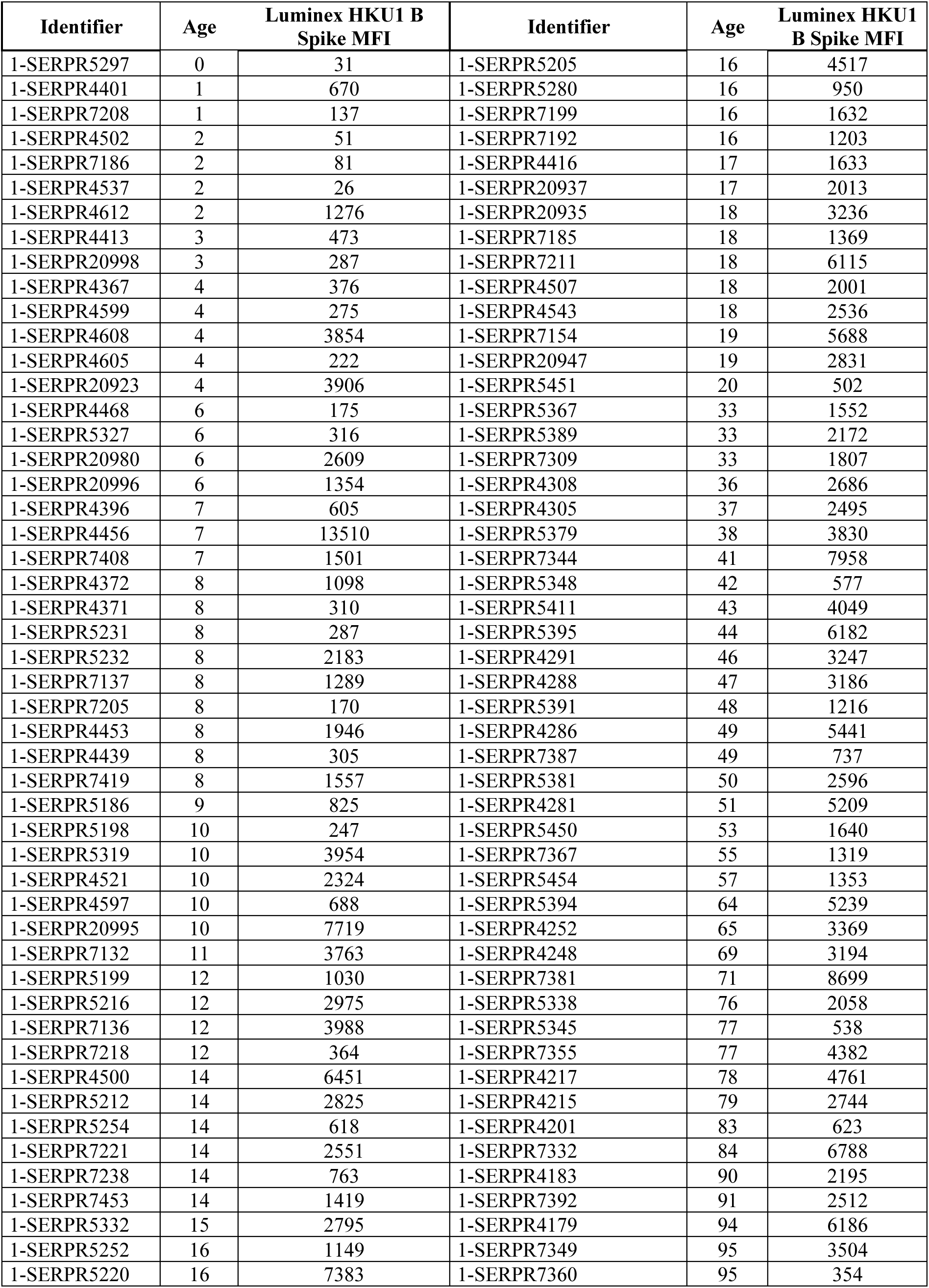
List of samples selected from SeroPed study ^8^ with sample identifier and associated HKU1 Spike luminex MFI measurement.

It has been reported that cross-reactive anti-coronavirus antibodies can be generated after SARS-CoV-2 infection or vaccination^52,53^. Whether these antibodies neutralize HKU1 has not been established. We used sera from a longitudinal cohort of 10 individuals sampled before and one month after administration of the Comirnaty^TM^ Omicron XBB.1.5 mRNA vaccine. Vaccination increased neutralization titers against SARS-CoV-2 D614G (12-fold) and XBB.1.5 (37-fold). However, it did not enhance anti-HKU1 A NAb titers and only slightly enhanced (1.4-fold) HKU1 B NAb titers (Figure 5c). The overall levels of anti-Spike antibodies, measured by flow cytometry^54^, were significantly increased for SARS-CoV-2 XBB.1.5 but not for HKU1 A and slightly (1.2-fold) for HKU1 B (Figure 5d).

## Discussion

The development of a robust culture system for HKU1 using Caco2-TMP2+ cells allowed us to study replication dynamics, viral entry and fusion, and host responses in cell lines and primary hAECs. We isolated two clinical viral strains, one HKU1 A and one HKU1 B that we further characterized. Viral stocks sequenced after 3 passages in Caco2-TMP2+ cells showed only minor stochastic variations. Both viruses grew exponentially in Caco2-TMP2+ cells, which formed syncytia upon infection and released infectious progeny. Virions displayed classical coronavirus morphology. Among a panel of cell lines expressing TMPRSS2, only Caco2-TMP2+ cells and to a lesser extent IGROV-1-TMP2+ and VeroE6-TMP2+ cells allowed potent infection, whereas others, such as A549-TMP2+ and MDCK-TMP2+ cells remained poorly permissive. This permissivity was not entirely explained by TMPRSS2 or sialic acid surface levels. Future work, exploring in these cell lines the conformation and activation of TMPRSS2, the O-acetylations on sialic acids and expression of other potential cofacotors will help determining the entry or post-entry replication steps that modulate HKU1 tropism.

Viral replication, both in engineered cell lines and in primary nasal hAECs, was profoundly influenced by temperature. HKU1 A and B replicated efficiently at 33°C, mimicking the upper airway environment, but were attenuated at 37°C. This is reminiscent of dynamics observed with other seasonal coronaviruses (NL63, OC43, and 229E)^27,28^ and common respiratory viruses (rhinovirus, RSV, and others) which replicate optimally at 32–34°C^34,35^, through mechanisms that are not fully understood. In primary hAECs, the IFN response was delayed at 33°C, suggesting that a slower triggering of antiviral cytokines and cellular restriction factors facilitates viral replication. Moreover, inhibiting the ISG response with Ruxolitinib partially rescued HKU1 replication at 37°C. Thus, there are probably both IFN-dependent and - independent mechanisms that impact HKU1 temperature sensitivity in hAECs.

The situation was somewhat different in Caco2-TMP2+ cells, in which the greater HKU1 replication at 33°C was apparently not directly linked to the IFN response. HKU1 did not trigger detectable ISGs in these cells and Ruxolitinib did not restore viral replication at 37°C. Moreover, HKU1 replicated more efficiently at 33°C than at 37°C in VeroE6-TMP2+ cells, which are defective for type 1 IFN response^55^. Therefore, in addition to delayed IFN-dependent innate immune responses, viral or cellular mechanisms mediate the temperature sensitivity of HKU1. We have identified some of these factors, but it is likely that others are involved. Spike-mediated cell-cell fusion was not greater but rather lower at 33°C than at 37°C, in line with reports with other coronavirus Spikes^29^. The block at higher temperature was not due to changes in TMPRSS2 surface expression, TMPRSS2 or HE enzymatic activity, or Spike surface levels, but was associated with reduced stability of cell-free virions and suppressed viral RNA synthesis 24 h after infection. This last point indicates that a late stage of the viral life cycle or that secondary infections are impacted at 37°C and deserves further investigation. It will be worth determining whether viral enzymes and proteins, the cellular environment or proviral factors may be sub-optimally active at 37°C. An alternatively, but not mutually exclusive explanation is that premature cell death of infected cells leading to abortive infection or some cellular antiviral mechanisms that are not triggered by IFN may be less effective at 33°C than at 37°C. It is important to note that while Caco2 cells are an interesting platform for HKU1 study, they are colorectal adenocarcinoma cells which physiologically never experience 33°C. Any cellular factor identified in Caco2 cells will thus need to be validated in primary cells.

Of note, HKU1 B was more temperature-sensitive than HKU1 A and barely replicated at 37°C in cell lines. Further experiments with additional viral isolates, including HKU1 C^10^, are needed to confirm this observation.

In contrast to HKU1, SARS-CoV-2 (KP3.1.1 variant) was not attenuated at 37°C. This confirms previous findings that SARS-CoV-2 replicate at both 33°C and 37°C in primary cells, with differences between variants^26,27,29,30^. These differences in temperature sensitivity represent important parameters regulating viral tropism and pathogenicity.

We report that IFITM1 and LY6E are potent inhibitors of HKU1 Spike-driven fusion and pseudovirus entry, establishing these plasma membrane-resident ISGs as core restrictions for HKU1. These ISGs are of interest as they inhibit SARS-CoV-2 infection and syncytia formation^40,56–58^. IFITM1 and LY6E were active against HKU1 at both 33°C and 37°C, indicating that the improved HKU1 replication at lower temperature is not due to a loss of activity of these antiviral factors. IFITM2 and IFITM3 were less or not efficient against HKU1, likely reflecting differences in subcellular localization. IFITM1 and LY6E locate to the plasma membrane^46,47^, which is in line with the entry route of HKU1^13^. The efficacy of IFITM1 in the presence of TMPRSS2 distinguishes HKU1 from SARS-CoV-2, which is no longer sensitive to this restriction factor when the serine protease is present^40^. This sets the stage for future exploration of innate immunity’s molecular determinants in coronavirus pathogenesis. One limitation is that we focused on overexpressed IFITMs. Future work will address the role of endogenous IFITMs, as reports have shown differences in their action against SARS-CoV-2^59–61^.

We also developed an infectious HKU1 neutralization assay to measure functional antibody activity in human cohorts. Sero-neutralization titers increased with age, paralleling epidemiological observations of widespread exposure and seroconversion in childhood. There was partial cross-neutralization between HKU1 A and B isolates, consistent with the relatively low (∼85%) identity between HKU1 A and B Spikes^10,13^. Vaccination with SARS-CoV-2 XBB.1.5 mRNA did not potently boost HKU1-neutralizing antibodies, highlighting the antigenic distance between HKU1 and pandemic coronaviruses^62^.

In summary, we established a comprehensive model to study HKU1, highlighting temperature-dependent viral replication and sensitivity to innate and humoral immune responses. Our findings provide novel insights into the viral and host determinants that shape coronavirus biology and disease.

## Acknowledgements

The OS lab is funded by Institut Pasteur, ANRS-MIE, the Vaccine Research Institute (VRI) (ANR-10-LABX-77), Labex IBEID (ANR-10-LABX-62-IBEID), the HERA projects DURABLE (grant 101102733), LEAPS (grant no 101094685) and European Vaccine Hub (EVH). The E.S.-L. laboratory is funded by Institut Pasteur, the INCEPTION program (Investissements d’Avenir grant ANR-16-CONV-0005), the Ixcore foundation for research, the French Government’s Labex IBEID (ANR-10-LABX-62-IBEID), the HERA Project DURABLE (grant no 101102733) and LEAPS (grant no 101094685). F.A.R acknowledges funding from Institut Pasteur, CNRS and by grants ANRS-MIE Project EMERGEN - ANRS 0149 - PRODEVAR and ANR-22-CE35-0004 BAT-CoV-ASIA. The NanoImaging Core at Institut Pasteur is acknowledged for support with sample preparation, image acquisition and analysis. We also thank O. Mayboroda from the NanoImaging Core for her help. The NanoImaging Core was created with the help of a grant from the French Government’s Investissements d’Avenir program (EQUIPEX CACSICE - Centre d’analyse de systèmes complexes dans les environnements complexes, ANR-11-EQPX-0008). This work has used the computational and storage services (Maestro cluster) provided by the IT department at Institut Pasteur. We thank N. Aulner, N. Mahtal, J. Fernandes and the UtechS Photonic BioImaging core facility (Institut Pasteur) and P. Charneau for help in lentiviral pseudotype preparation. Work with UtechS Photonic BioImaging is funded by grant no. ANR-10-INSB-04-01, no. ANR-24- INBS- 0005 FBI (BIOGEN) and Région Ile-de-France programme DIM1-Health. We thank all the staff members of the occupational health and medicine department of the Hospices Civils de Lyon (HCL), especially Pr Jean-Baptiste Fassier, who contributed to the sample collection. Human biological samples and associated data were obtained from NeuroBioTec (CRB HCL, Lyon France, Biobank BB-0033-00046). COVID SER cohort was funded by ANRS-MIE (Emergen study, grant ANRS-0154) and institutional grants from Fondation des HCL and HCL. The funders of this study had no role in study design, data collection, analysis and interpretation or writing of this Article.

## Contributions

The experimental strategy was designed by J.B., E.T., J.P., A.W., N.S., D.P., I.F. and O.S. Data acquisition and analysis were performed by J.B., E.T., J.P., A.W., N.S., D.P., F.G.-B., I.S., J.S., V.B., I.F., F.B., C.P., F.P., M.P., M.J.-G., V.M., N.C. and E.S.-L. Vital materials and expertise were provided by L.A., C.B., S.T.-A., M.-A.R.-W., D.D., A.F. and M.W. The manuscript was written by J.B. and O.S. and edited by J.B., N.S., T.B., M.-A.R.-W., A.F., M.W., D.P., F.A.R., E.S.-L., O.S. All authors read and approved the final version of the manuscript.

## Material and Methods

### Plasmids

phCMV-HKU1A (RefSeq: <u>YP_173238.1</u>), phCMV-HKU1B/C (UniProtKB/Swiss-Prot: <u>Q0ZME7.1</u>), pCSDest-TMPRSS2-S441A, pLV-TMPRSS2-Hygro and pLV-TMPRSS2-S441A-Hygro were previously described^13^. pQCXIP-Empty control, pQCXIP-IFITM1, pQCXIP-IFITM2 and pQCXIP-IFITM3 plasmids were previously described^40^. phCMV-SARS-CoV-2 Wuhan Spike was previously described^54^. phCMV-SARS-CoV-2 XBB.1.5 was previously described^63^. pQCXIP-BSR-GFP11 and pQCXIP-GFP1-10 were a kind gift from Yutaka Hata^64^ (Addgene plasmid no. 68716 and no. 68715). pCSDest-TMPRSS2 was a kind gift from Roger Reeves^65^ (Addgene plasmid no. 53887). pcDNA3.1-hACE2 was from Hyeryun Choe^66^ (Addgene plasmid no. 1786). pQCXIP-LY6E was generated by gateway cloning. For HE, A codon-optimized version of HKU1 HE (GenBank: AGT17757.1) and OC43 HE (RefSeq : YP_009555240.1) were ordered as a synthetic gene (GeneArt, Thermo Fisher Scientific) and was cloned into a phCMV backbone (GeneBank: AJ318514), by replacing the VSV-G gene.

### Cell lines

HEK-293T (293T), U2OS, Vero E6, A549, IGROV-I, MDCK, HeLa and Caco-2/TC7 (Caco2) were from the American Type Culture Collection and cultured in Dulbecco’s Modified Eagle’s Medium (DMEM) with 10% fetal bovine serum (FBS) and 1% penicillin/streptomycin (P/S), GFP-split cells were previously described^39^ and cultured with 1 µg/ml of puromycin (InvivoGen). Caco2-TMP2^KO^, Caco2-TMP2+ and Caco2-TMP^S441A^ were previously described^13^. TMPRSS2 expressing U2OS, Vero E6, A549, IGROV-I, MDCK and HeLa were generated by transduction as previously described^13^. Cells stably expressing TMPRSS2 were cultured with 100 µg/ml hygromycin. All cell lines were routinely screened for mycoplasma. Cells were authenticated by genotyping (Eurofins).

### Reagents

Nirmatrelvir was from (MedChemExpress; HY-138687). DMSO and Ruxolitinib phosphate (HY-50858) were from Fisher Scientific. Neuraminidase (Sialidase from Arthrobacter ureafaciens) was from Roche. IFN-β1b and IFN-λ1 were from ImmunoTools.

### Viral strains

HKU1 B (HCoV-HKU1/NDL/IPP01/2022), SARS-CoV-2 D614G (hCoV-19/France/GES-1973/2020, GISAID ID: EPI_ISL_414631) and XBB.1.5 (hCoV-19/France/IDF-HEGP-77-48-2232847922/2022, GISAID ID: EPI_ISL_16353849) were previously described^67,68,13^. SARS-CoV-2 KP.3.1.1 (hCoV-19/France/BRE-RELAB-IPP06405/2024, GISAID ID: EPI_ISL_19205879) was provided by the National Reference Centre for Respiratory Viruses, hosted by the Institut Pasteur and headed by Marie-Anne Rameix-Welti.

### Isolation of a new HKU1 A viral strain

HKU1 A nasopharyngeal swab material was obtained in spring 2023 at Ambroise Paré Hospital (Boulogne Billancourt, France). Material was collected in Transport Medium (EliTechGroup) and stored at −80 °C. Presence of HCoV-HKU1 was determined by BIOFIRE® RESPIRATORY PANEL 2.1 plus (Biomerieux). The HKU1 A strain (HCoV-HKU1/FRA/IPP02/2024) was isolated as previously described^13^, on ALI culture by the National Reference Centre for Respiratory Viruses, hosted by the Institut Pasteur and headed by Marie-Anne Rameix-Welti. Briefly, MucilAir, reconstructed human nasal epithelium cultures were kept at 33 °C under a 5% CO_2_ atmosphere. Mucus was removed from the apical side of the culture and cells were infected with nasal swabs diluted at a 1:4 ratio in Universal Transport Medium (Copan) for 2 h at 33°C. Viral input was removed, and cells were washed three times with 200 μl of HBSS (Gibco). The apical side was harvested twice every 24 h for 7 days by adding 200 μl HBSS for 10 min at 33°C, collecting the liquid on the apical side and diluting it (1:1) in Universal Transport Medium (Copan). The harvest was centrifuged at 1,000*g* for 5 min to remove debris, 50 µl was used for qPCR and sequencing, and the remaining samples were stored at −80 °C.

### HKU1 amplification on Caco2-TMP2+ cells

1.5x10^3^ Caco2-TMP2+ were seeded in a 96 well plate and cultured over night at 37°C under a 5% CO_2_ atmosphere. On day 0, media was changed to 100 ul of DMEM 2% FBS 1% P/S. Cells were infected with 5 μl of day 7 supernatant (Passage 0 or P0) of HKU1 A or B produced on hAEC culture (equivalent to approximatively 2x10^6^ viral RNA copies). Caco2-TMP2+ cells were cultured at 33°C for 9 to 11 days until a cytopathic effect was visible. Supernatant (P1) was harvested and centrifuged at 1,000 *g* for 5 min to remove debris. 25 μl of P1 supernatant was used to infect Caco2-TMP2+ cells seeded at 9x10^4^ cells/well in a 24 well plate in 300 μl of DMEM with 2% FBS, 1% P/S at 33°C. 300 μl of day 10 supernatant (P2) was used to infect a T75 flask of Caco2-TMP2+ cells seeded at 3.5x10^6^ cells/flask. P3 supernatant was harvested at 4 days pi, centrifuged at 3500 rpm for 45 min at 4°C and stored at −80°C. Further stocks (P4) were amplified from the P3 stock, titrated using TCID50 and sequenced.

### Sequencing of the HKU1 strain

RNA was extracted from the cell supernatant using the QIAamp Viral RNA Kit (Qiagen) following the manufacturer’s instructions. Extracted RNA was treated with Turbo DNase (Ambion) followed by ribosomal RNA depletion^69^. RNA was reverse-transcribed into double-stranded cDNA using random hexamers and libraries prepared using the Nextera XT kit (Illumina), before sequencing on an Illumina NextSeq500 (2 × 75 cycles). Raw sequence data (human reads removed) were deposited in the Sequence Read Archive (https://www.ncbi.nlm.nih.gov/sra) under BioProject ID PRJNA1329235. To determine the sequence of the acidic tandem repeat region of the genome, the region was separately amplified using external primers (HKU1_ATR_L1 5′-ATGAAGCAATGGCCTCTCGT-3′ and HKU1_ATR_R1 5′-CACAGAACGCAACCAACAGT-3′) before Sanger sequencing.

### Genome assembly

Adaptors and low-quality sequences of raw reads were removed using Trimmomatic v.0.39 (ref ^70^). We assembled the trimmed reads using megahit v.1.2.9 (ref ^71^) with default parameters. The contigs were queried against the NCBI non-redundant protein database using DIAMOND v.2.0.4 (ref ^72^) to look for potential contaminants in addition to the detected HKU1 genome. The Sanger data was used with the assembled contigs to generate the final HKU1 scaffold on which the trimmed reads were mapped to generate the final consensus (Figure S1B). The mapping data were visually checked to confirm the accuracy of the obtained genome using Geneious Prime 2023. The sequence of the isolated virus HCoV-HKU1/NDL/IPP02/2024 was deposited in GenBank, with accession number PX441742.

### Phylogenetic and recombination analysis

All available complete HKU1 genome sequence data and metadata were retrieved from the Bacterial and Viral Bioinformatics Resource Center^73^ (https://www.bv-brc.org/) in September 2025. Sequences were aligned using MAFFT v.7.467 (ref ^74^), and the alignment was checked for accuracy using Geneious Prime v2025.2.2 (https://www.geneious.com/). We used a combination of six methods implemented in RDP4 (ref ^75^) (RDP, GENECONV, MaxChi, Bootscan, SisScan and 3SEQ) to detect potential recombination events in the newly reported genome. Multiple recombination signals were detected in the novel genome, but none was unique to this virus, and instead were congruent with the patterns observed with publicly available genomes falling in the same lineage. With these limitation in mind, the ModelFinder application^76^, as implemented in IQ-TREE v.2.0.6 (ref ^77^), was next used to select the best-fitting nucleotide substitution model, and maximum-likelihood phylogenies were inferred using complete genomes or the Spike coding sequences. Branch support was calculated using ultrafast bootstrap approximation with 1,000 replicates^78^. The phylogenies were visualized using the auspice module from Nextstrain^79^. Interactive phylogenies are available at https://github.com/Simon-LoriereLab/HKU1.

### 50% tissue culture infectious dose (TCID50) quantification

Culture supernatants were thawed and serially diluted (10-fold) from 10^−1^ to 10^−8^ in DMEM. Six replicates of viral dilutions (50 µL each) were seeded in flat bottom 96-well plates and mixed with 15.000 Caco2-TMP2^+^ cells in DMEM-3% FBS (150 µL).

Two readout methods were used to calculate the viral titer. The first method used immunostaining: after 72 h at 33°C, cells were fixed in 4% PFA for 30 min at room temperature (RT) and stained with pan coronavirus antibody mAb10 and Hoechst (1:10,000). Plates were imaged on an Opera Phenix High-Content Screening System (Revvity). The second method used TCID50 calculation. Cells were fixed 6 days pi and living cells were stained with crystal violet. Titer was determined as the lowest viral concentration without positive staining in 50% of the six replicates.

### Sample preparation for electron microscopy

HKU1 A and HKU1 B samples, corresponding to supernatants from infected cells showing strong cytopathic effects, were clarified by centrifugation for 30 minutes at 3.500 rpm. Viral supernatants were fixed in 4% PFA for 30 min at room temperature. HEPES was added to achieve a 25 mM final concentration, followed by filtration through a 0.45 μm filter. Fixed supernatant was gently layered onto a 20% sucrose cushion, prepared in Tris buffer (50 mM Tris, 1 mM EDTA, 100 mM NaCl) in an ultra-clear tube (344059 Dutscher) and ultracentrifuged at 30.000 rpm for 90 min at 4°C. The pellet was resuspended in 10 μl of Tris buffer. Samples were analyzed by Transmission electron microscopy (TEM) and cryo-electron microscopy.

### Transmission electron microscopy (TEM)

Standard 200 mesh formvar coated and carbon stabilized grids were discharged for 15s with 2 mA at 0.1 mBar using a Quorum Q150 ES unit. 3 µl of sample was pipetted onto the activated grid and incubated for 3 minutes at RT before the grid was washed three times in Mili-Q water for 30 seconds respectively and finally stained with 2% uranyl acetate for 1 minute at RT. After staining, grids were dried using Whatman paper. Negative stain images were acquired using a TEM Tecnai T12 BioTWIN 120 kV with a Rio16 detector.

### Grid preparation for cryo-electron microscopy

Grids were prepared with HKU1 A viral particles isolated by ultracentrifugation on a sucrose cushion and mixed with 5 nm Protein A-Au beads. 3 μl of the sample were applied to Lacey carbon films on 300 mesh copper grids (Agar scientific) that had been glow-discharged using a Pelco glow discharge system at 15 mA for 25 s. Samples were immediately vitrified in 100% liquid ethane using a Mark IV Vitrobot (Thermo Fisher Scientific) by blotting for 3 s after a 15 s wait with Whatman no. 1 filter paper at 8°C and 100% relative humidity.

### Tilt series data collection

Cryo-ET datasets were collected using the Tomo-EPU automated image acquisition software (Thermo Fisher Scientific) on a 300 kV cold-Field Emission Gun Titan Krios transmission electron microscope (Thermo Fisher Scientific) equipped with a Falcon 4i direct electron detection camera (Thermo Fisher Scientific) and a Selectris X energy filter (Thermo Fisher Scientific). For tilt series acquisition the microscope was set in nanoprobe mode, energy filter at 20 eV (zero loss) and an objective aperture of 100. Tilt series were acquired with a dose dose-symmetric acquisition scheme^80^ ranging from +60° to −60° with 3° increments at the nominal magnification of 64,000 corresponding to a pixel size of 2.0 Å, with defocus ranging from -2 to -4 µm. The total dose applied was 140 e−/Å^2^ with an initial electron dose of 3.5 e−/Å^2^ at 0 angle and an increased exposure time function of the tilt angle (4.0 e−/Å^2^ at ±60°) to partially compensate for the ice thickness increase.

### Cryo-ET data analysis

Fifty tilt series were recorded as eer (electron-event representation) files. Twenty of them, containing at least one virus and without recording artefact, were processed on a Linux workstation, using a set of shell scripts calling MotionCor3^81^, IMOD^82^ routines and standard Unix tools. MotionCor3 was used to convert the eer frames in 6 fractions (corresponding to a dose of ∼0.5e^-^/Å^2^/fraction), to perform the gain correction (gain file flipping along the x axis) and to align the fractions (3x3 patches, 15 iterations, tolerance of 0.2). The coherence between the tilt angle values and the defocus gradient of the images was checked by determining the defocus value of each image (IMOD *ctfplotter*) with and without inverting the tilt angles (with and without *-invert* flag). The stacks were built (*newstack*) and, accordingly to the results of the previous verification and to previous tests showing that the handedness needed to be changed when using eer files in our setup, flipped along the tilt (i.e. x) axis (*clip flipx*). They were manually checked, and the bad images were removed. For each resulting stack, the images were coarsely aligned (*tiltxcorr*), the fiducials were identified (*autofidseed*), tracked (*beadtrack*), and the alignment parameters determined (*tiltalign*). The alignment quality was checked, by looking at the residual error local mean and the values of the largest residual (*tiltalign* log file), and the fiducial tracking was verified when the residual error local mean was found larger than 1 pixel (2 Å) and manually corrected when necessary (*3dmod*). The aligned stack was generated with a binning factor of 2 resulting in a final pixel size of 4 Å. It was ctf-corrected using the defocus values previously determined (*ctfplotter*). The received and cumulated electron doses corresponding to each image were extracted from the mdoc file and the images were dose-weighted (*mtffilter*). Finally, two tomograms were built (*tilt; clip rotx*) by using weighted back projection, one with a sirt-like filter equivalent to 8 iterations, one without filtering, after manual adjustment of the tomogram center and thickness and of the intensity dynamic: we first calculated each tomogram by using a final *thickness* of 500 planes (200 nm) and a *shiftz* of 0. We also set the *scale* parameter to a “reasonable” value (in our case a scaling of 20 for the sirt-like filtered tomogram and 5 for the non-filtered tomogram). We then adjusted the *thickness* and *shiftz* so that the viruses are entirely contained within the tomograms, with the smallest possible thickness, and we modified the *scale* parameters to ensure that the minimal and maximal values of the final map are -15 000 and +15 000.

For each tomogram, we manually determined the virus particle locations. As the viruses have a globally prolate shape, we approximate their diameter and rotation axis length in *3dmod*. The virus particle volume was calculated by adapting the formulae 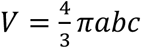 (where a,b and c are the radius along the three ellipsoid axes) and 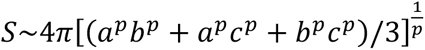 (where p=log_2_(3)=1.6075, Knud Thomsen approximation^83^) and the values were rounded to two significant digit as the local deformation will likely prevent to give more precise values. The number of Spikes at the surface of each virus particle was measured by template matching (pytom^84^ version 1.1) following by manual curation (*3dmod*): the reference was built from the coordinates of the pdb3jcl.ent file in a box of 100^3^ voxels with a pixel size of 4 Å, inverted, and filtered at 5 nm resolution. Template matching (pytom_match_template.py) was performed with a spherical max, a particle diameter of 200 Å, a symmetry of 3 and a low pass filtering of 50 Å) and restricted in a box of ∼550x550x300 pixels around each virus. It resulted in a new map assigning a score value to each pixel. The 100 best matches were extracted from this map (pytom_extract_candidates.py with an exclusion radius of 25 pixels), ordered along the z axis coordinates and converted to an IMOD model (*point2model*). The model was superimposed with the tomogram in *3dmod* and all candidate points were checked. The ones that were not corresponding to a spike protein were removed. We also verified that they were no false negative: i.e. all visible spike proteins were detected by pytom with the parameters we used.

### Western blot

HKU1 A and HKU1 B samples were clarified by centrifugation for 30 minutes at 3.500 rpm. Clarified supernatants were gently layered onto a 20% sucrose cushion, prepared in Tris buffer in an ultra-clear tube (344059 Dutscher) and ultracentrifuged at 30.000 rpm for 90 min at 4°C. Viral particles were lysed in 1% Triton X-100 and 1X Roche cOmplete protease inhibitors for 30 min on ice. Equal amounts of lysates were separated by sodium dodecyl sulfate-polyacrylamide gel electrophoresis (SDS-PAGE) and transferred onto a nitrocellulose membrane. Membranes were stained with the following antibodies diluted in western blot buffer (PBS, 1% BSA, 0.05% Tween, 0.01% Na Azide): Rabbit anti-HKU1 S1 Polyclonal Antibody (Thermo Fisher Scientific, PA5-120768, 1:2000) and Rabbit anti-HKU1 S2 Polyclonal Antibody (Thermo Fisher Scientific, PA5-120769, 1:1000) and a polyclonal sera (G4) from the SeroPed cohort^8^. Species-specific fluorescent secondary antibodies were used. Fluorescent signals were detected with a LI-COR Odyssey scanner and analyzed using ImageStudioLite software.

### HKU1 infection in cell lines

Caco2-TMP2^KO^, Caco2-parental, Caco2-TMP2+, Caco2-TMP2^S441A^ and the following parental and TMPRSS2 overexpressing cell lines: U2OS, Vero E6, A549, IGROV-I, MDCK and HeLa were seeded in a 96 well plate at 10^4^ cells/well. Cells were infected with a range of MOIs (calculated on Caco-TMP2+ cells) of HKU1 A and B in DMEM 10% FBS 1% P/S and kept at 33°C under a 5% CO_2_ atmosphere. Cells were fixed 4 days post infection in 4% PFA for 20 min and stained with mAb10 at 1 μg/ml for 1 h at RT followed by anti-human Alexa Fluor 488-or 647-conjugated Goat anti-Human Antibody (Thermo Fisher Scientific, 1:500) for 1 h at RT and then by Hoechst at 1:10.000 for 10 min. Images were acquired on an Opera Phenix High-Content Screening System (Revvity) and the Spike positive area was quantified using SignalImageArtist High-Content Imaging and Analysis Software.

### Flow cytometry

Surface expression of TMPRSS2 was assessed on live cell lines by staining with anti-TMPRSS2 VHH A01-Fc^13^ at 1 μg/ml for 30 min at 4 °C in MACS buffer, followed by staining with Alexa Fluor 647-conjugated Goat anti-Human Antibody (Thermo Fisher Scientific, A-21445, 1:500). The results were acquired using an Attune Nxt Flow Cytometer (Life Technologies).

For surface receptor expression quantification, MucilAir^TM^, reconstructed human nasal epithelium cultures that had been differentiated at air-liquid interface for four weeks from pooled and single donors, were purchased from Epithelix. Cells were cultured at 33°C or 37°C for 72 h. Inserts were washed once with PBS+EDTA 0.1% and cells were dissociated by adding 1 ml of TrypLE™ Express (12605-010) in the basal chamber and 200 ul in the apical chamber and incubating the cells for 10 min at 37°C. Cells were washed and expression level of ACE2, TMPRSS2, aminopeptidase-N (APN or CD13) and dipeptidyl peptidase-4 (DPP-4 or CD26) was assessed on live cells by surface staining in MACS buffer with anti-ACE2 VHH B07-Fc^85^ or anti-TMPRSS2 VHH A01-Fc at 1 μg/ml, CD13-PE (130-120-312, Miltenyi Biotec, 1:50) and CD26-PE (130-126-41, Miltenyi Biotec, 1:50) for 30 min at 4 °C. For VHHs, a secondary antibody anti-human Alexa Fluor 488- or 647-conjugated Goat anti-Human Antibody (Thermo Fisher Scientific, 1:500) was used. The cells were washed twice with PBS and fixed with 4% PFA. For intracellular staining, fixed cells were stained with Rabbit anti-β-IV-tubulin-AF488 (ab204003, Abcam, 1 μl/ml) and Rabbit anti-cytokeratin V AF647 (ab193895, Abcam, 0.1 μl/ml) for 30 min at room temperature in PBS 1% BSA 0.01% NaAz. The results were acquired using an Attune Nxt Flow Cytometer (Life Technologies). Analysis was performed with FlowJo 10.10.0.

Transfection efficiency for 293T fusion experiments was assessed on live cells for TMPRSS2, ACE2 and MHC-I and on fixed cells for IFITMs and Spike. For TMPRSS2, ACE2 VHH A01-Fc and VHH B07-Fc were used as described above at 1 μg/ml. For MHC-I, cells were stained with HLA-ABC Monoclonal Antibody (W6/32, eBioscience™), followed by secondary GAM 647. For the Spike, transfection efficiency was measured at the surface of live cells using mAb10 diluted in MACS buffer for 30 min at 4 °C and a human secondary IgG. mAb10 is an antibody generated from a SARS-CoV-2 infected patient that cross-reacts with HKU1^86^. For IFITMs and Spike, cells were fixed 4% PFA, washed and stained intracellularly with anti-IFITM1 (#60074-1-Ig, Proteintech) or anti IFITM2/3 (#66081-1-Ig, Proteintech) at 1:250 in PBS/BSA/Azide/0.05% Saponin. The results were acquired using an Attune Nxt Flow Cytometer (Life Technologies).

### Sialic acid staining

Different cell lines, cultured under a 5% CO_2_ atmosphere, were seeded at 2x10^5^ cells/well in 24 well plates and treated for 24h with 50 mU/ml neuraminidase from *Arthrobacter ureafaciens* (Roch). Cells were harvested using Trypsin/0.1% EDTA, washed in PBS and stained 1 h at 4 °C in PBS/1% SVF containing either 10 µg ml−1 *Sambucus nigra agglutinin* Lectin-FITC (Lectin NSA) (Thermo Fisher Scientific, L32479) or 2.5 µg ml−1 recombinant mouse Siglec-E-Fc (BioLegend, 551504). Cells were washed twice in PBS. Lectin-stained cells were fixed for 10 min in 4% paraformaldehyde. mSiglec-E-Fc stained cells were further incubated with Alexa Fluor 647-conjugated Goat anti-Human Antibody (Thermo Fisher Scientific, A-21445, 1:500) for 30 min at 4 °C before being fixed for 10 min in 4% paraformaldehyde. Data were acquired on an Attune NxT instrument (Life Technologies).

### HE activity

HEK293T cells were transfected in suspension using Lipofectamine 2000 (ThermoFisher), with 50 ng of HKU1/OC43 HE or HKU1 S, adjusted to 100 ng DNA with pQCXIP-Empty and incubated 24 h at 37°C. 24 h post-transfection, cells were equilibrated to 33°C or 37°C for 30 min prior to the assay. Fresh 4-Nitrophenyl acetate (pNPA; MERCK, N8130) was prepared by adding stock solution in pre-warmed (33° or 37°C) serum-free DMEM. Cell culture media was then replaced with 90 µL of a serial dilution of pNPA substrate. After 15 min incubation at the respective temperatures, 405 nm absorbance was measured using an EnSpire^TM^ 2300 plate reader (Revvity) maintained at the corresponding temperature (33°C or 37°C).

### Human nasal epithelium temperature sensitivity experiments

MucilAir^TM^, reconstructed human nasal epithelium cultures that had been differentiated for four weeks from pooled donors, were purchased from Epithelix and cultured in 700 μl MucilAir media on the basal side of the air/liquid interface cultures and monitored for healthy cilia movements. Cultures were kept at 37°C under a 5% CO_2_ atmosphere. Three days before infection, basal media was changed to 700 μl MucilAir media containing either DMSO (1:4500 dilution) or 10 μM Ruxolitinib and placed at either 33°C or 37°C under a 5% CO_2_ atmosphere. One hour before infection, mucus was removed from the apical side of the culture by washing with Hanks’ Balanced Salt Solution (HBSS) (Gibco). Basal media was changed to new media containing the respective treatment. ALI cultures were infected in 200 μl MucilAir media containing HKU1 A (P4) at 1:200 dilution for 2 h at 33 or 37°C. Viral input was removed, and cells were washed three times with 200 μl of HBSS (Gibco). The apical side was harvested every 24 h for 4 days by adding 200 μl HBSS for 10 min, collecting the liquid on the apical side. The harvest was centrifuged at 1,000 *g* for 5 min to remove debris, 50 µl was used for qPCR, and 75 µl was used for Luminex multiplex assay, and the remaining samples were stored at −80 °C. ALI cultures were then washed with HBSS and fixed for 30 min in 4% PFA for subsequent immunofluorescence staining.

### Analysis of HKU1 replication by real-time qPCR

Viral RNA was extracted from culture supernatants using the Quick-DNA/RNA Viral 96 Kit (Zymo) according to manufacturer’s instructions. qPCR was run using the Luna Universal Probe One-Step kit on a QuantStudio 6 (Thermo Fisher Scientific) according to manufacturer’s instructions using the following primers from ref^3^: HKUqPCR5, 5′-CTGGTACGATTTTGCCTCAA-3′ and HKUqPCR3, 5′-ATTATTGGGTCCACGTGATTG-3′ and TaqMan QSY probe 5′-FAM-TTGAAGGCTCAGGAAGGTCTGCTTCTAA-QSY7-3′). A DNAString (ref^13^) was used to generate a standard curve (5′-GGATCCTACTATTCAAGAAGCTATCCCTACTAGGTTTTCGCCTGGTACGATTTTGCCTCAAGGCTATTATGTTGAAGGCTCAGGAAGGTCTGCTTCTAATAGCCGGCCAGG TTCACGTTCTCAATCACGTGGACCCAATAATCGTTCATTAAGTAGAAGTAATTCTAATTTTAGACATTCTGATTCTATAGTGAAACCTG-3′).

### Luminex multiplex assay

ALI supernatants were inactivated with 1% Triton X100 (vol/vol) for 2 h at room temperature. Samples were prepared according to manufacturer’s instructions and acquired using the Luminex Intelliflex instrument. Cytokine concentrations were calculated from MFIs using theProcartaPlex application (https://apps.thermofisher.com/apps/procartaplex/#./). Samples with concentrations below the limit of detection of the assay were assigned a value of half the lowest detectable cytokine.

### Immunoflorescence staining

Tissues on multiple ALI inserts were permeabilized with 0.5% Triton X-100 for 20 minutes at RT. Blocking was performed with PBS containing 20% FBS and 0.3M glycine for 1 h at RT. Intracellular staining was performed overnight at 4°C using primary antibodies in PBS with 1% BSA and azide. The primary antibodies used were: mAb10 at 1 μg/ml, anti-MX1 at 1:100, and VHH-TMPRSS2 at 6 μg/ml. Afterward, the ALI inserts were washed three times with PBS for 5 minutes each. Secondary antibody incubation was carried out for 3 h at RT, with Hoechst at 1:10,000 for nuclear staining. Imaging was performed on a Leica Thunder at 10x magnification for Spike quantification and on a Leica SP8 confocal microscope at 40x magnification for MX1 and TMPRSS2 quantification.

### Image quantification

For Spike quantification the entirety of the stained insert (1/4^th^) was imaged at 10x magnification using identical settings and computational clearing. In ImageJ, for each image the following steps were applied, first background was removed, second a fix threshold, calculated on a negative sample, was applied, objects smaller than a cell were removed and the Spike positive area (above threshold) was calculated. Percentage positivity was calculated by dividing by the total cell area. For MX1 and TMPRSS2 quantification, for each sample, Z-stack images of four different regions were acquired on Leica SP8 confocal microscope using identical settings. For analysis, maximum Z-projects were generated, and mean fluorescence intensity of each image was calculated.

### ISG quantification in ALI cultures at 33°C and 37°C

MucilAir^TM^, reconstructed human nasal epithelium kept at 37°C were placed at 33°C or 37°C for 72 h. Inserts were washed with PBS+EDTA 0.1% and cells were dissociated by adding 1 ml of TrypLE™ Express (12605-010) in the basal chamber and 200 μl in the apical chamber and incubating the cells for 10 min at 37°C. Cells were washed with PBS, pelleted and lysed using RNA lysis buffer (QIAGEN). RNA extraction was performed using the RNeasy Plus Mini Kit (QIAGEN) according to the manufacturer’s instructions. PCR was run using the Luna Universal One-Step kit on a QuantStudio 6 (Thermo Fisher Scientific) according to manufacturer’s instructions.

### ISG quantification in Caco2-TMP2+ cells cultured at 33°C and 37°C

Caco2-TMP2+ were seeded in a 12 well plate at 10^5^ cells/well in DMEM 10% SVF 1%P/S alone or containing 1000 U/mL IFN-β1b and placed at either 33°C or 37°C. After 2 h30, 6 h and 24 h; cells were washed with PBS+EDTA 0.1% then lysed with RLT buffer (QIAGEN) and stored at -80°. RNA extraction was performed using the RNeasy Plus Mini Kit (QIAGEN) according to the manufacturer’s instructions. PCR was run using the Luna Universal One-Step kit on a QuantStudio 6 (Thermo Fisher Scientific) according to manufacturer’s instructions.

### ISG quantification in infected and bystander Caco2-TMP2+ cells after IFN-β1b stimulation

Caco2-TMP2+ were seeded in a 96-well plate (μClear, no. 655090) for 16h at 37°C. Cells were infected with HKU1 A or B at indicated MOIs and incubated for 24h 33°C. Cells were wash with PBS and fresh media containing nothing (Non-treated) or 1000 U/mL of IFN-β1b and placed at either 33°C or 37°C for an extra 24h. Cells were fixed 4% PFA and stained intracellularly with anti-S mAb10 (0,1 μg/ml), anti-IFITM1 (#60074-1-Ig, Proteintech) or anti IFITM2/3 (#66081-1-Ig, Proteintech) at 1:250 in PBS/BSA/Azide/0.05% Saponin followed by anti-human Alexa Fluor 488- or 647-conjugated Goat anti-Human, Goat-anti-Mouse or Goat-anti-Rabbit Antibodies (Thermo Fisher Scientific, 1:500) for 1 h at RT and then by Hoechst at 1:10,000 for 10 min. Images were acquired on an Opera Phenix High-Content Screening System (Revvity). Spike positive and negative area was delimited and mean fluorescence intensity of IFITM1 and IFITM2/3 staining in S+ and S- areas was quantified using SignalImageArtist High-Content Imaging and Analysis Software.

### Primers

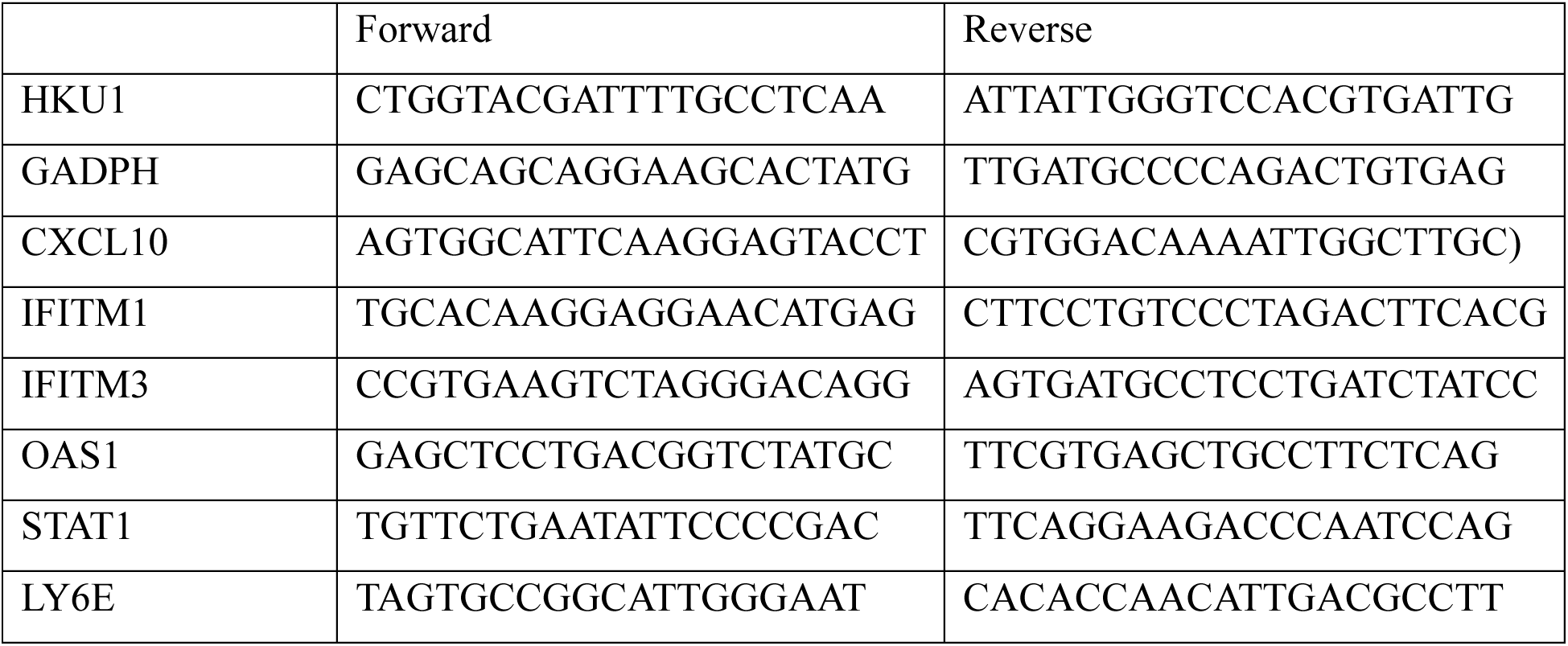

### GFP-split fusion assay

Cell–cell fusion assays were performed as previously described^39,40^. Briefly, 293T cells stably expressing GFP1-10 and GFP11 were cocultured at a 1:1 ratio (6 × 10^4^ cells per well) and transfected in suspension with Lipofectamine 2000 (Thermo Fisher Scientific) in a 96-well plate (μClear, no. 655090) (10 ng of Spike plasmid, 20 ng IFITM1, IFITM2, IFITM3 or LY6E plasmid and 8 ng of TMPRSS2 plasmid adjusted to 100 ng DNA with pQCXIP-Empty). Cells were imaged 24 h post transfection.

For acceptor–donor experiments, 293T GFP-split cells (GFP1-10 and GFP11) were transfected separately with 1000 ng of DNA (150 ng Spike, 250 ng TMPRSS2 plasmid adjusted to 1000 ng with pQCXIP-Empty) and plated in a 12 well plate and placed at 37°C. 24h post transfection cells detached in PBS+EDTA, washed and acceptor and donor cells were mixed and seeded at 6 × 10^4^ cells per well. Cells were imaged 2 h 30 post co-culture. For all experiments, images covering 90% of the well surface were acquired on an Opera Phenix High-Content Screening System (Revvity). The GFP area was quantified on SignalImageArtist High-Content Imaging and Analysis Software.

### Lentiviral pseudovirus generation

Pseudoviruses were produced by transfection of 293T cells as previously described^87^. Briefly, cells were co-transfected with plasmids encoding for lentiviral proteins, a luciferase reporter and the HKU1 Spike plasmid. Pseudotyped virions were harvested two and three days after transfection. Production efficacy was assessed by measuring infectivity or HIV Gag p24 concentration.

### Pseudovirus infection

293T cells were transfected in suspension using Lipofectamine 2000 (20 ng ISG plasmids, 8 ng TMPRSS2 plasmids, adjusted to 100 ng DNA with pQCXIP-Empty) and incubated 24 h at 37°C. The next day, infection was performed with spinoculation (2 h at 1200G) using 2 μL of pseudoviral production per well. After spinoculation, 100 μL of media was added and cells were incubated at 33°C or 37°C. After 24 h, 150 μL of media was carefully removed and 50 μL of Bright-Glo lysis buffer (ProMega) was added. After 6 min, luminescence was acquired using the Enspire in a white plate (Revvity).

### TMPRSS2 enzymatic activity

Enzymatic activity was measured using Boc-QAR-AMC (R&D Systems, ES014), a substrate of TMPRSS2 that fluoresces when cleaved. 75 μL of DMEM medium containing 3.75 ng/µl or 6 ng/µl of soluble TMPRSS2^13^ was placed in a black 96 well plate and incubated at 33°C or 37°C for 30 min. 25 µl of cold PBS containing 100 µM of fluorogenic substrate was added and fluorescence was assessed using the previously heated Enspire for 1h at 33°C or 37°C, with an acquisition occurring every 2 min.

### IFN response in Caco2-TMP2+ cells

Caco2-TMP2+ cells were seeded in a 96 well plate at 10^4^ cells/well in 100 µL DMEM 10%FBS 1% P/S containing either 100 µM of Ruxolitinib or DMSO. The day after, cells were infected with HKU1 at a MOI of 0.1 for 2 h or treated with a range of IFN concentration. Cells were fixed 1-day pi in 4% PFA for 20min and stained with mAb10 and anti-IFITMs at 1 μg/ml for 1 h at RT followed by anti-human Alexa Fluor 488- or 647-conjugated Goat anti-Human Antibody (Thermo Fisher Scientific, 1:500) for 1 h at RT and then by Hoechst at 1:10,000 for 10 min. Images were acquired on an Opera Phenix High-Content Screening System (Revvity) and the Spike positive area was quantified using SignalImageArtist High-Content Imaging and Analysis Software.

### Effect of drugs on HKU1 infection

Caco2-TMP2+ cells were plated at 10^4^ cells/well in DMEM 10% FBS, 1% P/S, in a 96 well plate and treated with a range of concentrations of Neuraminidase, IFN-β1b or IFN-λ and placed at 37°C under a 5% CO_2_ atmosphere. For Nirmatrelvir conditions, cells were treated 2 h pre-infection. At 24h post treatment, cells were infected in DMEM 3% FBS 1% P/S containing the relevant treatments with HKU1 A or B at MOI 0.01, SARS-CoV-2 KP.3.1.1 at a dilution of 1:600, D614G at a dilution of 1:10,000 and BA.1 at a dilution of 1:500. Cells were fixed for 30 min in 4% PFA 48h post-infection. PFA was removed and HKU1 infected cells were stained for 45 min at RT in PBS/BSA/Azide/0.05% Saponin with primary mAb10 at 0.5 µg/mL. SARS-CoV-2 infected cells were stained with human monoclonal antibody NCP-1^88^ at 0.5 µg/mL. Primary antibody staining was followed by secondary Alexa Fluor 488-conjugated Goat anti-Human Antibody (Thermo Fisher Scientific, 1:500).

### Effect of VHHs on HKU1 infection

Caco2-TMP2+ or Caco2-TMP2^S441A^ cells were plated at 10^4^ cells/well in DMEM 10% FBS, 1% P/S, in a 96 well plate and maintained at 37°C under a 5% CO_2_ atmosphere overnight. 2 h pre-infection, cells were treated with a range of concentrations of monomeric VHH-A01, VHH-A07 and non-target control VHH-93 in DMEM 3% FBS 1% P/S. Cells were infected with HKU1 A or B at MOI 0.01, or SARS-CoV-2 KP.3.1.1 at a dilution of 1:600. Cells were fixed for 30 min in 4% PFA 48h post-infection. PFA was removed and HKU1 infected cells were stained for 45 min at RT in PBS/BSA/Azide/0.05% Saponin with primary mAb10 at 0.5 ug/mL and SARS-CoV-2 variants infected cells were stained with human monoclonal antibody NCP-1^88^ at 0.5 µg/mL. Primary antibody staining was followed by secondary Alexa Fluor 488-conjugated Goat anti-Human Antibody (Thermo Fisher Scientific, 1:500).

### Effect of temperature on HKU1 replication in Caco2 cells

Caco-TMP2+ and Caco-TMP^KO^ cells were seeded in 48-well plates at 60,000 cells per well in DMEM 10% FBS, 1% P/S and incubated for 24 h at either 33°C or 37°C under a 5% CO_2_ atmosphere. On Day 1, cell lysis and supernatant collection were performed prior to infection with HKU1 A and B. Infections were carried out in 120 ul of DMEM with 3% FBS for 2.5 h at 33°C or 37°C, cells were washed 2 times for 5 min and once for 10 min with DMEM 3% FBS followed by incubation in fresh medium at the specified temperature. Samples were collected at various time points pi (2.5, 4, 6, 8, 10, 24, and 48 h) for cellular and viral RNA extraction, followed by RT-qPCR analysis. For cellular RNA, extraction was performed using the RNeasy Plus Mini Kit (QIAGEN) according to the manufacturer’s instructions. For viral release, supernatant was collected, and RNA was extracted using Quick-DNA/RNA Viral Kit (Zymo) according to manufacturer’s instructions. PCR was run using the Luna Universal One-Step kit on a QuantStudio 6 (Thermo Fisher Scientific) according to manufacturer’s instructions. For cellular RNA, primers for GAPDH and HKU1 were used to analyze expression by 2^-ΔΔCt^. For viral release, HKU1 signal was quantified by normalizing to a standard curve.

### S-Flow assay

293T cells were transfected with SARS-CoV-2 or HKU1 phCMV-Spike or a control plasmid (pQCXIP-Empty) using Lipofectamine 2000 (Life Technologies). One day after, transfected cells were detached using PBS-EDTA and transferred into U-bottom 96-well plates (50,000 cells per well). Cells were incubated at 4°C for 30 min with sera (1:300 dilution) in PBS containing 0.5% BSA and 2 mM EDTA, washed with PBS, and stained using anti-human IgG Fc Alexa Fluor 647 antibody (109-605-170, Jackson Immuno Research). Cells were washed with PBS and fixed for 10 min using 4% PFA. Data were acquired on an Attune NxT instrument (Life Technologies). Specific binding was calculated with the following formula: 100 × (% binding on 293T-S − binding on control cells)/(100 − binding on control cells).

### Neutralization assay

For HKU1, Caco2-TMP2+ cells were plated at 2×10^4^ cells per well in a μClear 96-well plate (Greiner Bio-One). The indicated HKU1 strain was incubated with serially diluted sera for 15 min at RT and added to the cells. The sera were heat inactivated 30 min at 56 °C before use. 24 h post infection, cells were fixed with 4% PFA, washed and stained with mAb10-AF488 at 2 µg/ml and Hoechst (1:1,000 dilution; Invitrogen). Images were acquired with an Opera Phenix high-content confocal microscope (Revvity). The Spike positive area and the number of nuclei were quantified using the Harmony software (Revvity). The percentage of neutralization was calculated using the area of Spike as the value with the following formula: 100 × (1 − (value with serum − value in ‘noninfected’)/(value in ‘no serum’ − value in ‘noninfected’)). Neutralizing activity of each sera was expressed as the ED_50_. ED5_0_ values (in dilution values) were calculated using a reconstructed curve with the percentage of neutralization at the different indicated concentrations.

For SARS-CoV-2, S-Fuse assay was used as previously described^89^. U2OS-ACE2 GFP1–10 or GFP 11 cells, also termed S-Fuse cells, become GFP^+^ cells when they are productively infected with SARS-CoV-2. Cells were mixed (at a 1:1 ratio) and plated at 8×10^3^ cells per well in a μClear 96-well plate (Greiner Bio-One). The indicated SARS-CoV-2 strains were incubated with sera at the indicated dilutions for 15 min at room temperature and added to S-Fuse cells. Sera were heat inactivated 30 min at 56 °C before use. 18 h pi, cells were fixed with 2% PFA, washed and stained with Hoechst (1:1,000 dilution; Invitrogen). Images were acquired with an Opera Phenix high-content confocal microscope (Revvity). The GFP area and the number of nuclei were quantified using the Harmony software (Revvity). The percentage of neutralization was calculated as previously described^89^. Neutralizing activity of each sera was expressed as the ED_50_.

### Effect of temperature on virion stability

HKU1 A or B virus preparations were incubated at 4°C, RT (about 22°C), 33°C and 37°C. Every hour, samples at respective temperatures were placed on ice. At 4 h post incubation, each viral dilution was used to infect Caco2-TMP2+ cells, with each condition tested in triplicate to ensure data reliability. Caco2-TMP2+ cells were placed at 33°C under a 5% CO_2_ atmosphere.

24 h post-infection, cells were fixed in 4% PFA for 20 min, washed PBS and stained intracellularly in PBS/BSA/Azide/0.05% Saponin with mAb10-AF488 at 2 μg/ml. Hoechst (1:1,000 dilution; Invitrogen) was added before plate acquisition on an Opera Phenix High-Content Screening System (Revvity) and the Spike positive area was quantified using Harmony High-Content Imaging and Analysis Software (Revvity).

### SeroPed Cross-Sectional Study

Sample selection for this study was performed using the SeroPed database^8^. From this database, samples from participants of all ages were considered. We organized these samples by ascending HKU1-Spike antibody dilution to capture the assay’s full dynamic range. From this ordered list, we selected 100 samples, ensuring they represent the spectrum of antibody responses.

## Ethical statement

Samples from the SeroPed study were leftovers from routine medical blood sample processing in French hospital laboratories, processed in accordance with existing regulations and guidelines of the French Commission for Data Protection (Commission Nationale de l’Informatique et des Libertés). Sera were completely anonymous, and it was not possible to return to individual patients’ files.

For the Covid-Ser study, written informed consent was obtained from all participants. Ethics approval was obtained from the national review board for biomedical research in April 2020 (Comité de Protection des Personnes Sud Méditerranée I, Marseille, France; ID RCB 2020-A00932-37), and the study was registered at ClinicalTrials.gov (NCT04341142).

## Supplemental Figures

**Supplemental Figure 1.**
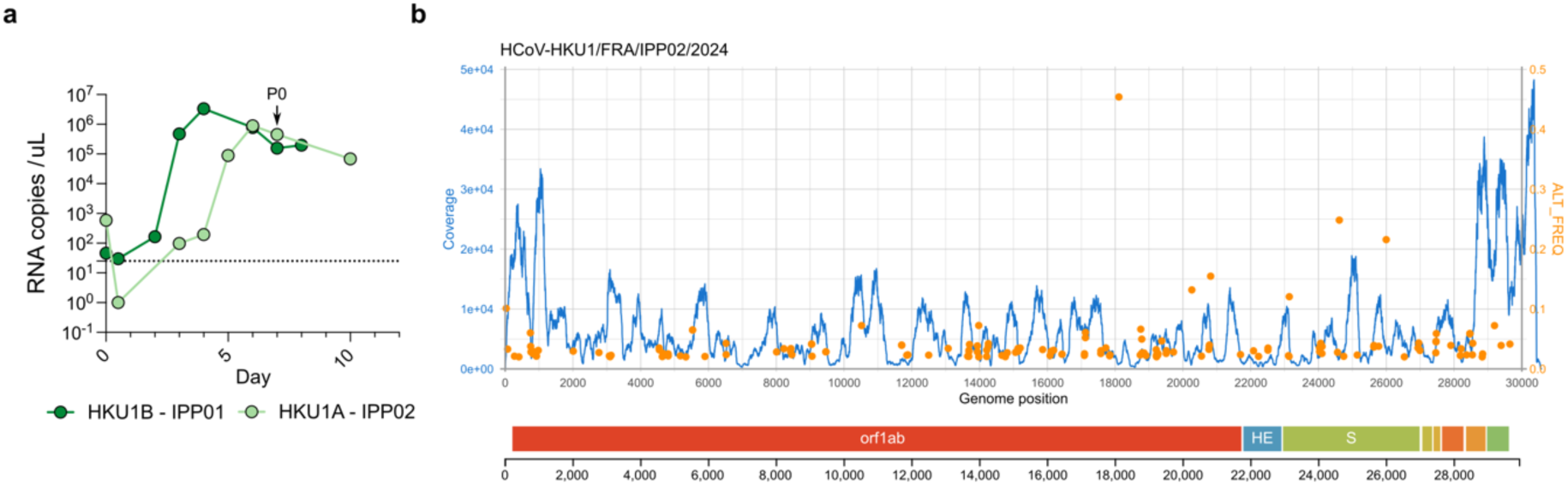
a Isolation of new HKU1 isolates. Two HKU1 clinical isolates were amplified on primary hAECs at 33°C as described^1313^. Viral RNA copies in the supernatant of hAECs cells were quantified by RT-qPCR. The values at day 0 represent the viral input. P0 marks the supernatant used for further viral amplification on Caco2-TMP2+ cells. **b Reads coverage and frequency of minor variants of HCoV-HKU1/FRA/IPP02/2024 (Virus passaged 3 times on Caco2-TMP2+ cells)**. Intra-sample single-nucleotide variants frequencies were estimated using iVAR^90^. The genome organization of HKU1 is shown below the plot.

**Supplemental Figure 2.**
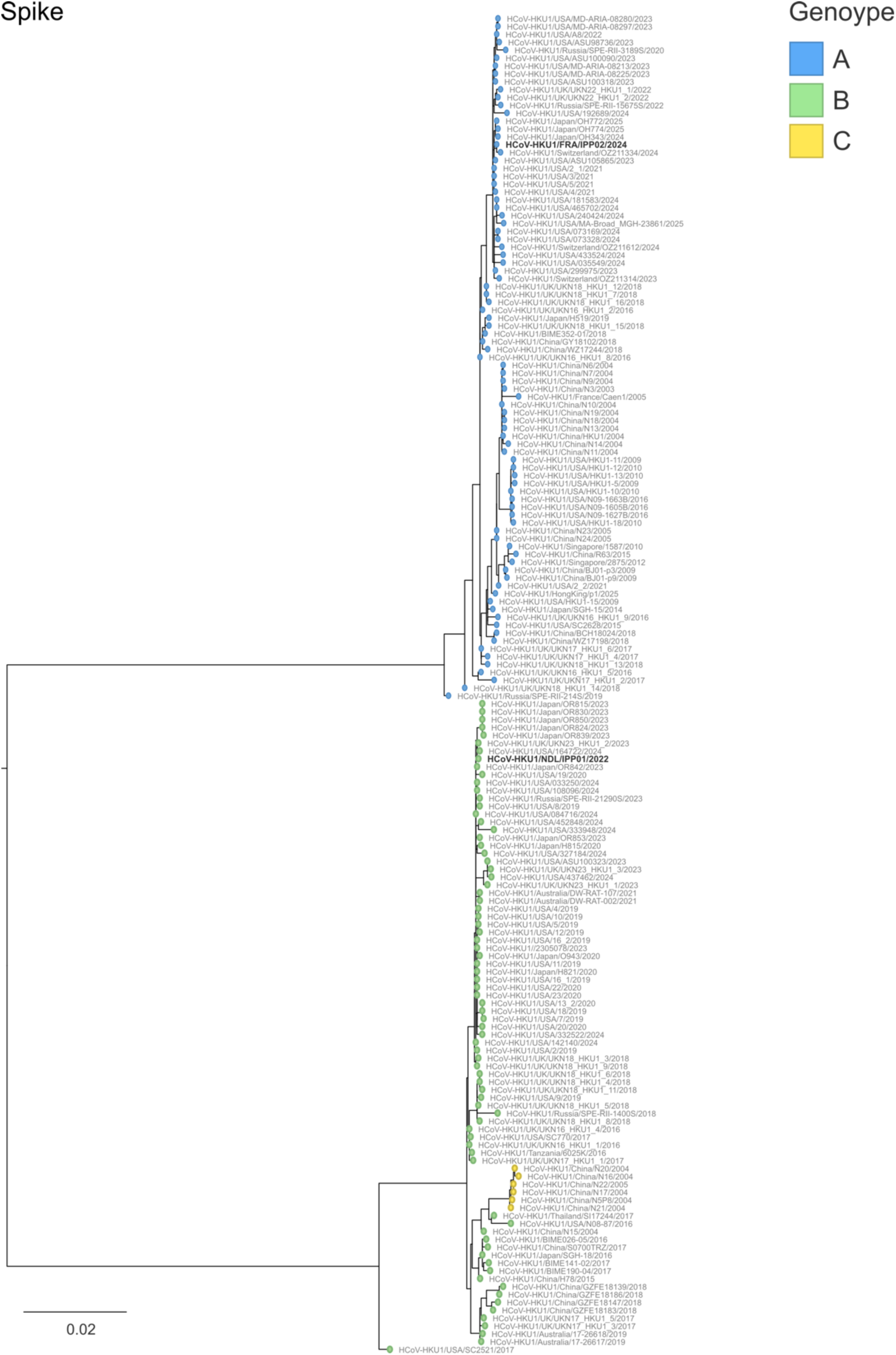
Complete genome phylogenetic analysis of HKU1. Maximum likelihood phylogenies of human coronavirus HKU1 (n = 184) estimated using IQ-TREE v2 with 1000 replicates from the complete viral genome sequences. The tree is midpoint rooted and ultrafast bootstraps values are shown on the main branches. The tip name corresponding to the isolates used in this study are highlighted in bold and underlined.

**Supplemental Figure 3.**
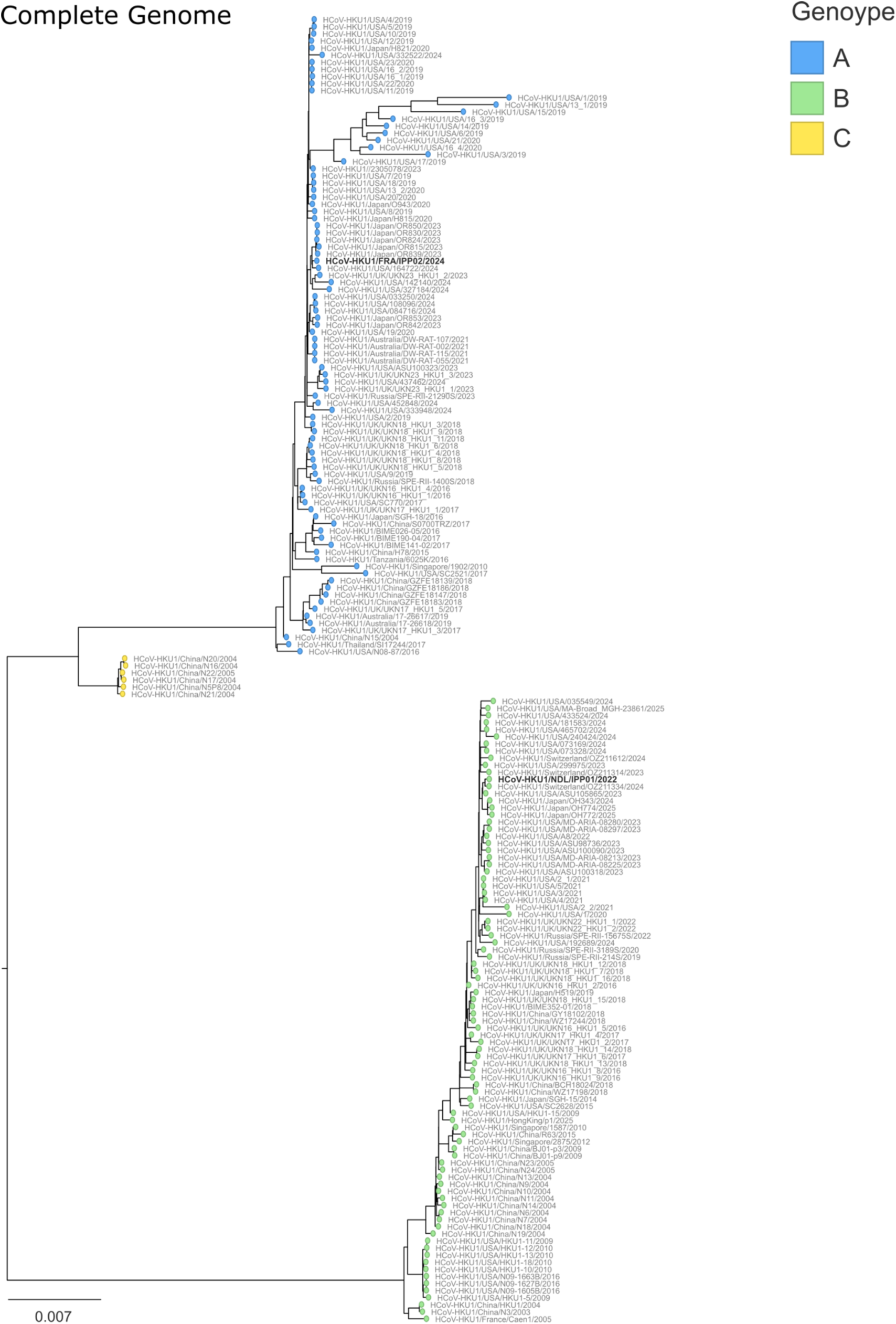
Spike phylogenetic analysis of HKU1. Maximum likelihood phylogenies of human coronavirus HKU1 (n = 184) estimated as in Figure S2 from the Spike coding sequences. The tip name corresponding to the isolates used in this study are highlighted in bold and underlined.

**Supplemental Figure 4.**
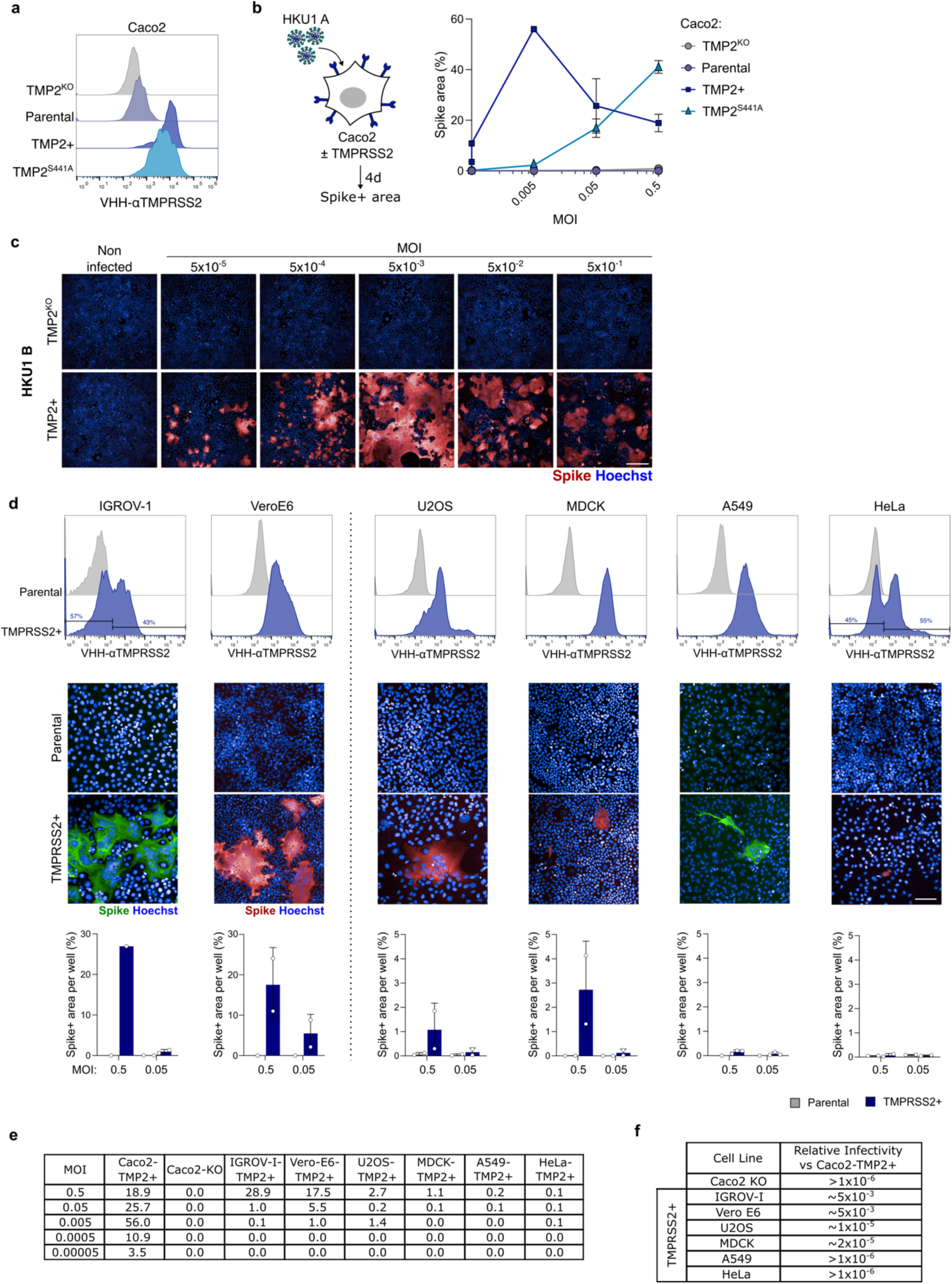
a Surface levels of TMPRSS2 in Caco2 cells stably expressing WT or catalytically inactive TMPRSS2^S441A^. Cells were stained for TMPRSS2 using VHH-A01-Fc and analyzed by flow cytometry. Representative histograms of Caco2-TMP2^KO^ (Light grey), unmodified parental cell line (Blue), Caco2-TMP2+ (Dark blue) or Caco2-TMP2^S441A^ cells (Light blue). **b Caco2-TMP2+ cells are permissive to HKU1.** Various Caco2 cell lines were infected at the indicated MOI with HKU1 A and analyzed four days post infection. Spike area of images (as represented in Figure 1g) was quantified. **c HKU1 B showed similar infectivity to HKU1 A in Caco2-TMP2+ cells.** Representative images of Caco2-TMP2+ cells infected with HKU1 B at indicated MOIs. Spike (red), Hoechst (Blue). Scale bar = 300 μm. **d TMPRSS2 expression and representative images of cell lines of Figure 1H**. (Top) Surface levels of TMPRSS2 in parental and TMPRSS2-transduced cell lines. Cells were stained for TMPRSS2 using VHH-A01-Fc and analyzed by flow cytometry. Representative histograms of parental (light gray) and stably TMPRSS2-transduced (dark blue) cell lines are shown. (Middle) Representative images of parental and TMPRSS2 transduced cell lines infected with HKU1 A at MOI 0.5. Spike (red or Green), Hoechst (Blue). (Bottom) Spike area at different MOIs was quantified. Data are mean ± SD of 1 to 3 independent experiments. Scale bar = 100 μm. **e** Exact values of S area for each cell line tested (heatmap Fig. 1h). **f Approximate permissivity of various cell lines overexpressing TMPRSS2 compared to Caco2-TMP2+ cells.** The ratio between lowest MOI triggering infection in Caco2-TMP2+ cells compared to that in the respective cell line is indicated.

**Supplemental Figure 5.**
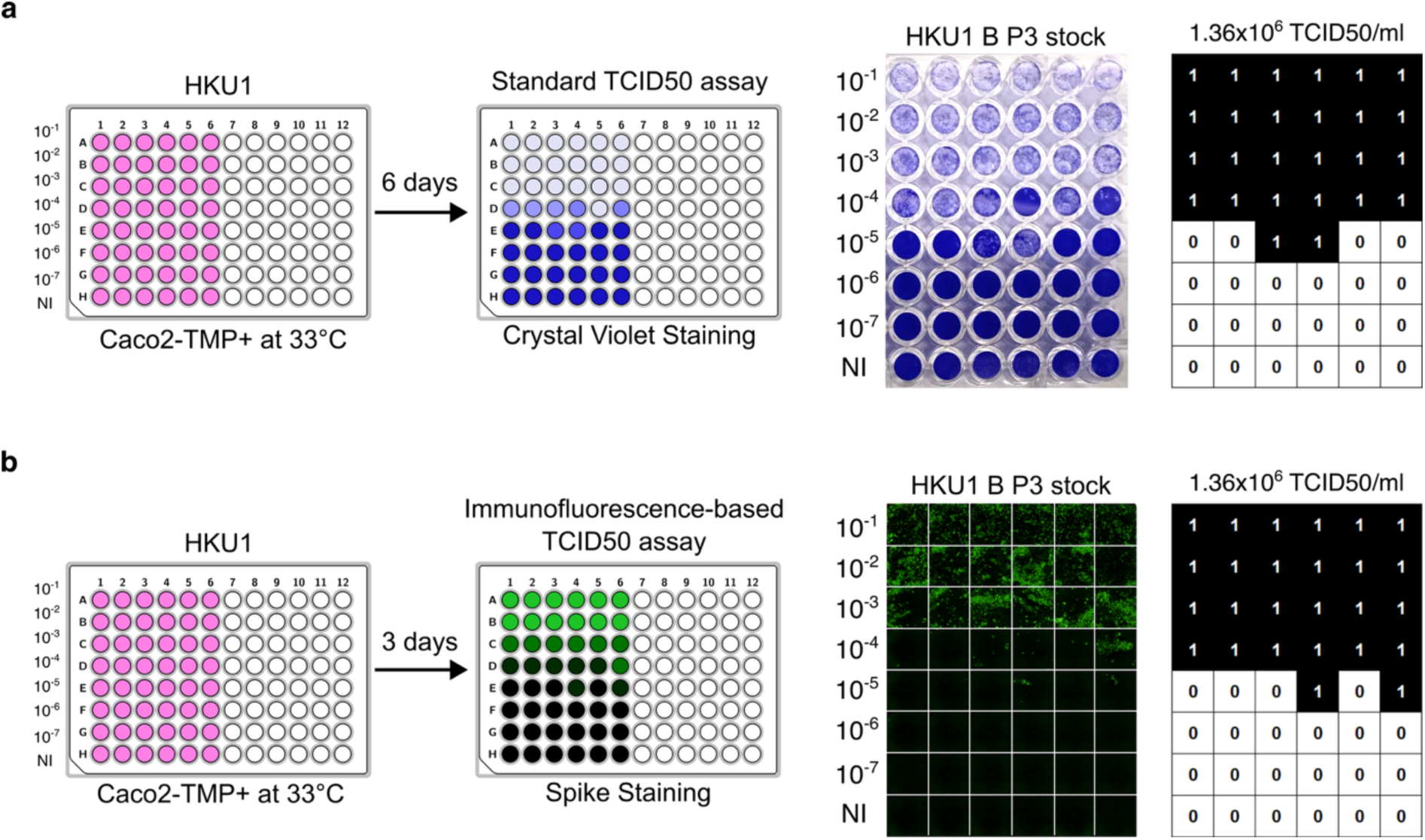
Titration of HKU1 on Caco2-TMP2+. Caco2-TMP2+ were plated at confluence in low serum media at 33°C and infected with serial dilutions of a HKU1 stock. **a** Six days post infection, plates were stained with crystal violet and cytopathic effect (ECP) was quantified by eye. **b** Three days post infection, plates were stained with anti-S antibody and imaged. Positive wells were identified by S staining. TCID50 was calculated as described in ref ^91^.

**Supplemental Figure 6.**
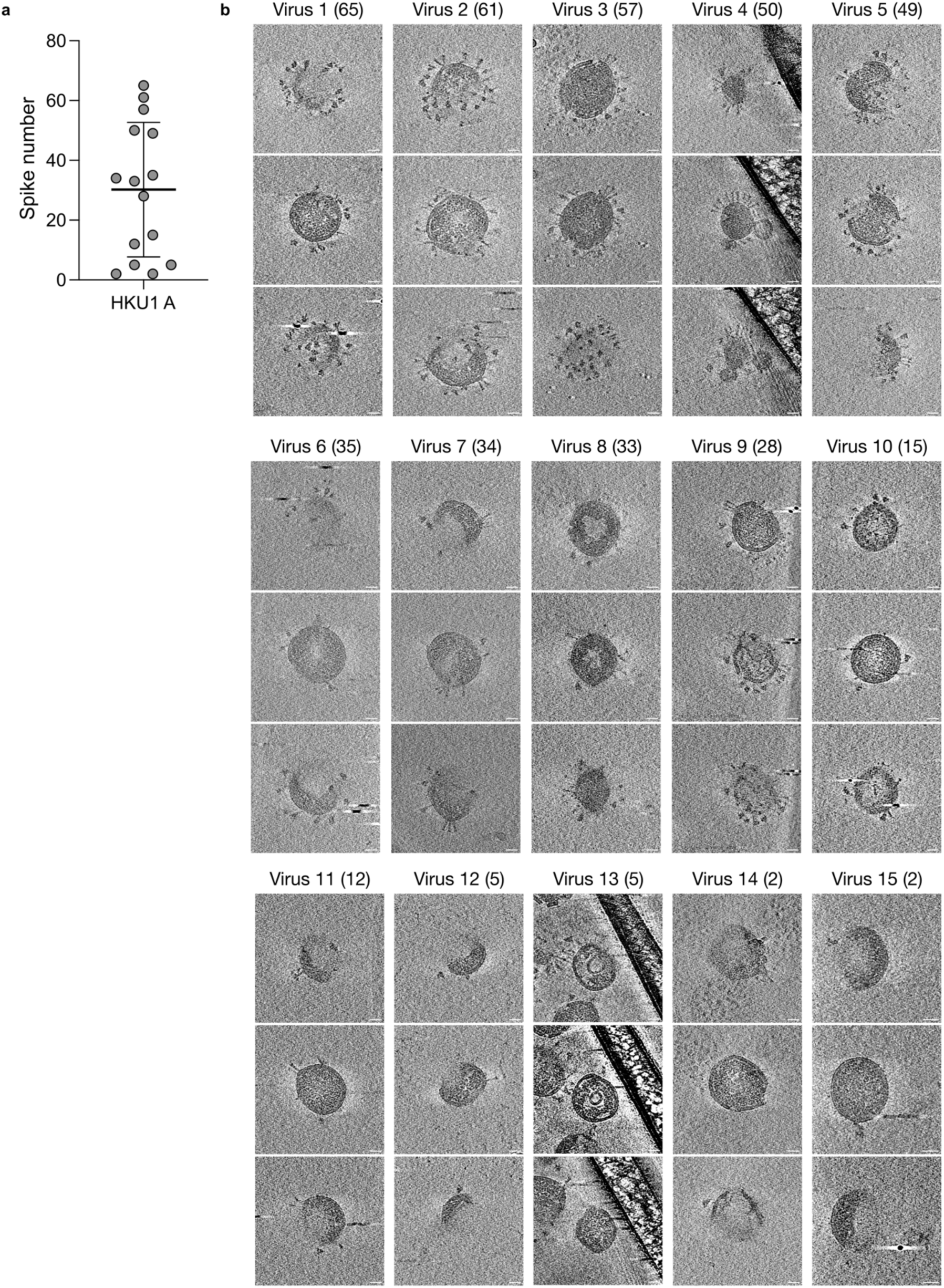
Cryo-electron tomogram of HKU1 A particles produced in Caco2-TMP+ cells and purified by ultracentrifugation. **a** Number of Spikes per particle. **b** Cryo-electron tomograms. Three images are shown per viral particle. Distinct surface Spike glycoproteins are visible, projecting from the viral envelope. Scale bar = 25 nm.

**Supplemental Figure 7.**
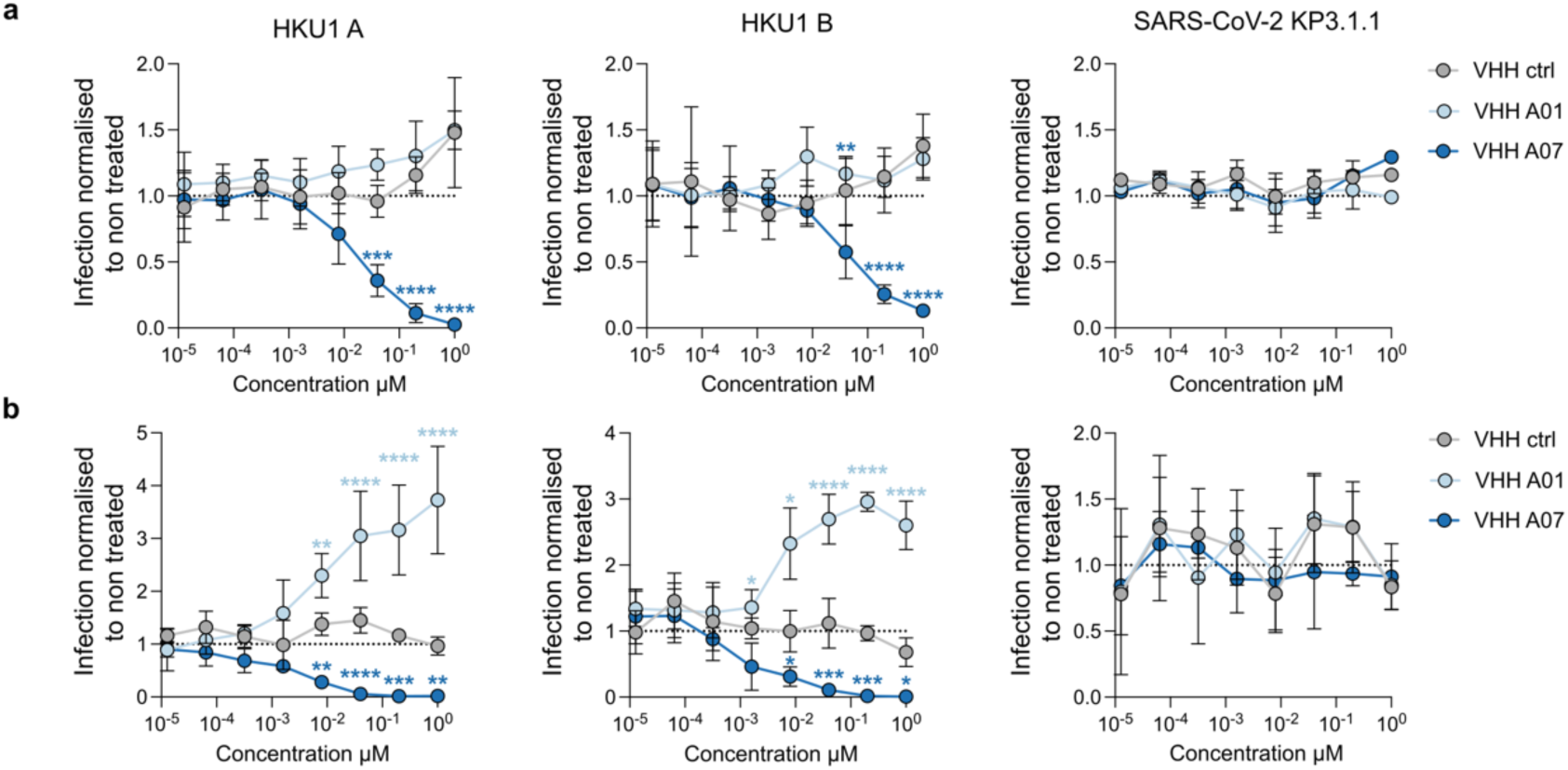
Anti-TMPRSS2 VHHs inhibit infection of Caco2-TMP2+ or Caco2-TMP2^S441A^ cells. Caco2-TMP+ (**a**) or -TMP2^441A^ cells (**b**) were pretreated with VHH-ctrl (non-targeting), VHH A01 (Binds TMPRSS2 but does not block HKU1 Spike-TMPRSS2 interaction) and VHH A07 (Binds TMPRSS2 and blocks HKU1 Spike-TMPRSS2 interaction) infected with HKU1 A/B or SARS-CoV-2 (KP.3.1.1 variant). Infection was measured after 48 h by Spike area quantification. Infection was normalized to the non-treated condition. Data are mean ± SD of 4 independent experiments. Statistical analysis: Two-way ANOVA Dunnett’s multiple comparisons test compared to non-target VHH *p<0.05, **p<0.01, ***p<0.001, ****p<0.0001.

**Supplemental Figure 8.**
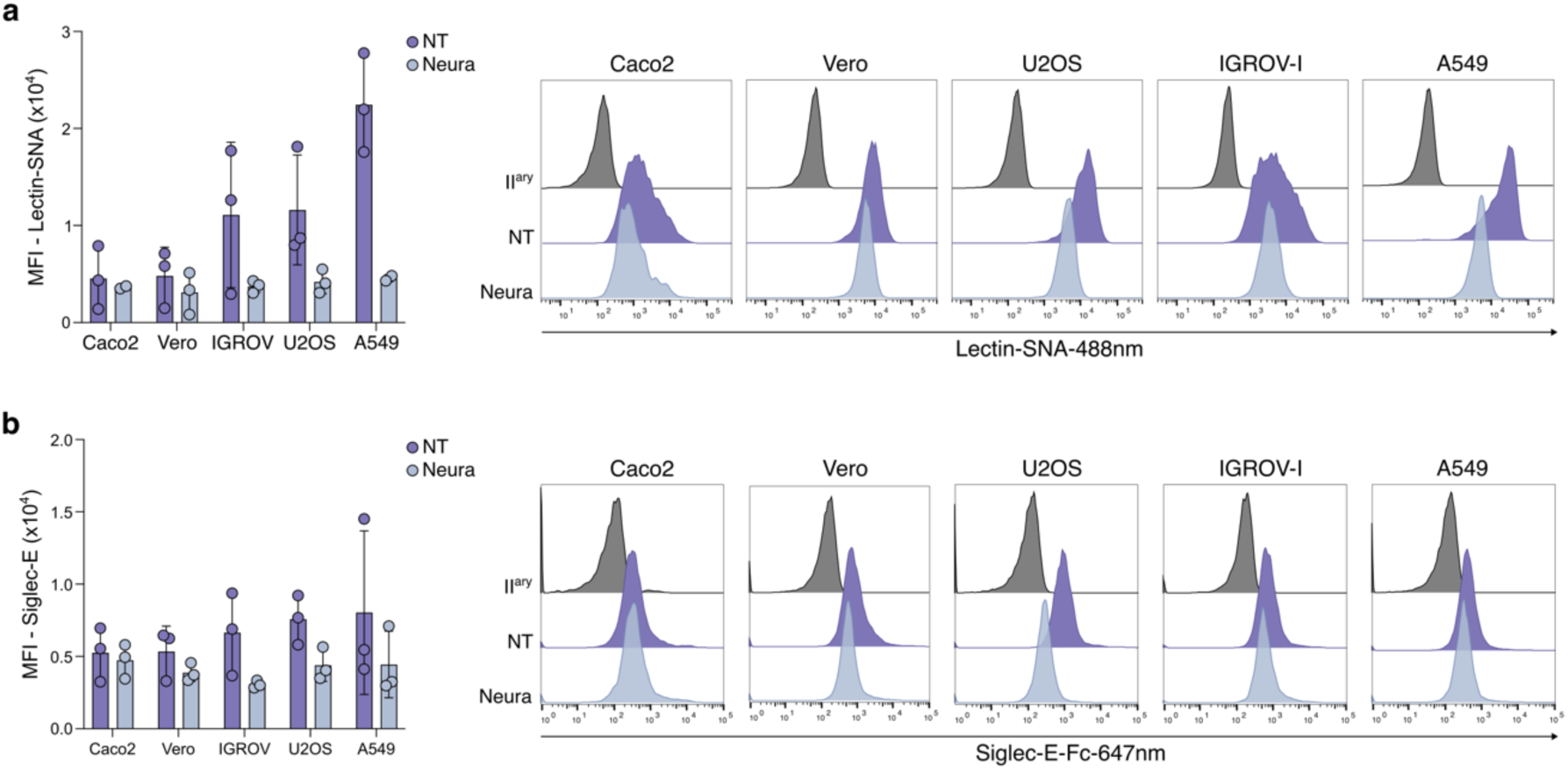
Sialic acid expression levels on cell lines. Indicated cell lines were treated for 24h with Neuraminidase at 50 mU/ml and stained with **a** SNA lectin and **b** mSiglec-E. Left: Mean fluorescence intensity quantification. Right: representative histograms. Data are mean ± SD of 3 independent experiments.

**Supplemental Figure 9.**
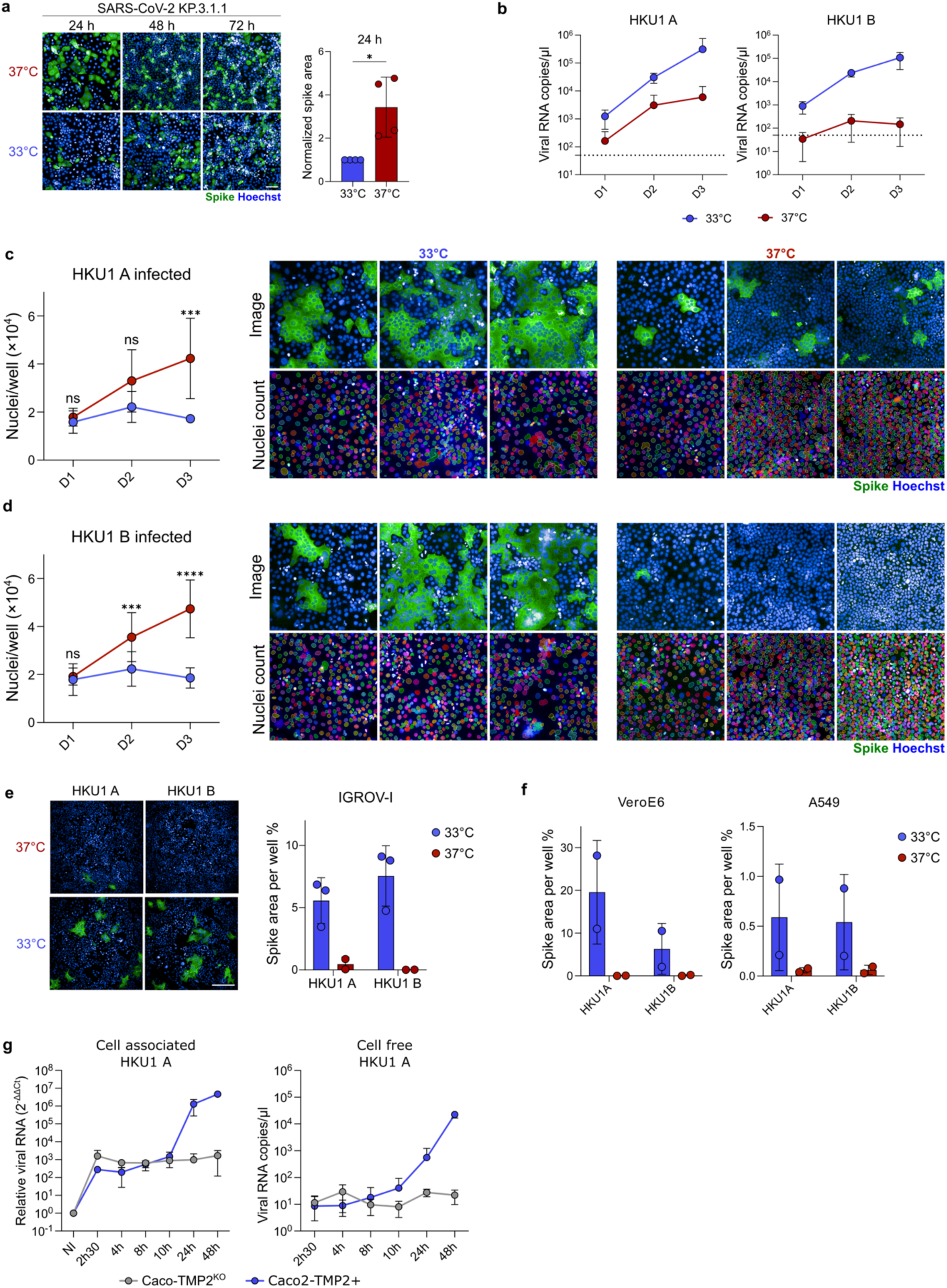
a SARS-CoV-2 (KP.3.1.1 variant) replicates more efficiently at 37°C than at 33°C in Caco2-TMP2+ cells. Caco2-TMP2+ were infected with SARS-CoV-2 KP.3.1.1 and kept at 33°C or 37°C for three days. Cells were fixed daily for Spike staining. (Left) Representative images and Spike area quantification 24h pi. Scale bar = 100 μm. (Right) Spike area quantification. Data are mean ± SD of 4 independent experiments. Statistical analysis: Unpaired t test with Welch’s correction. *p<0.05. **b** Caco2-TMP2+ cells were infected with HKU1 (MOI 0.1) at 33°C or 37°C for three days. Supernatants were collected for quantification of viral genomes over time by qPCR at 33°C and 37°C. Data are presented as viral copies/ul and are mean ± SD of 2 independent experiments. **c-d Lack of substantial cell death of HKU1 infected caco2-TMP2+ at 37°C.** Number of nuclei per well over time in HKU1 A (**c**) or HKU1 B (**d**) infected cells at 33°C and 37°C. Left: Nuclei/well quantification. Right: representative images of Hoechst staining and nuclei delimitation. Data are mean ± SD of 4 independent experiments, with 9 analyzed wells per experiment. Statistical analysis: Two-way ANOVA with Šídák’s multiple comparisons test. **e-f Impact of temperature on HKU1 replication in various cell lines. e** IGROV-1-TMP2+ (Figure S5d) were infected with HKU1 A or B at MOI 0.5, stained for Spike at 2 days pi. Representative images (left) and Spike area quantification (right) are shown. Data are mean ± SD of 2-3 independent experiments. Scale bar = 300 μm. **f** VeroE6-TMP2+ and A549-TMP2+ cells, (Figure S5d) were infected with HKU1 A or B at MOI 0.5 and stained for Spike at 4 days pi. Spike area quantification are shown. Data are mean ± SD of 2 independent experiments. **g Cell-free and cell-associated viral RNA over time at 33°C and 37°C in Caco2-TMP2^KO^ cells.** Caco2-TMP2^KO^ were infected and processed as described in figure 2f.

**Supplemental Figure 10.**
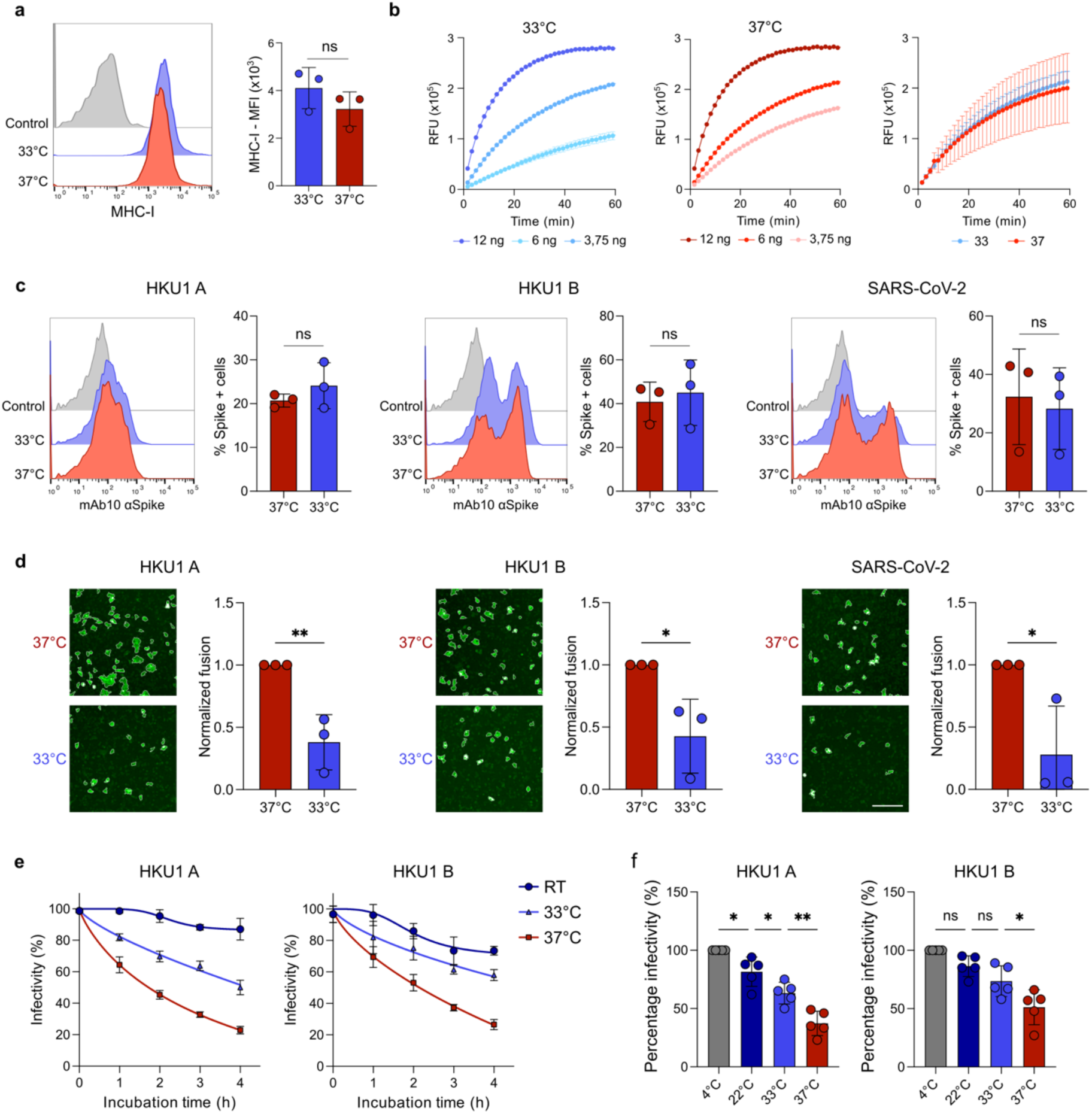
Effect of temperature on TMPRSS2 activity, Spike expression and fusion, and virion stability. **a MHC-I expression at 33°C and 37°C.** Caco2-TMP2+ cells cultured for 48 h at 33°C or 37°C were stained for MHC-I and analyzed by flow cytometry. (Left) Representative histograms of MHC-I expression of Caco2-TMP2+ cultured at 33°C (Blue) and 37°C (Red) and cells stained with secondary antibody alone (Gray). (Right) Quantification of MHC-I MFI. Data are mean ± SD of 3 independent experiments. Statistical analysis: Unpaired t-test. ns = non-significant. **b Enzymatic activity of TMPRSS2 at different temperatures.** Catalytic activity of indicated quantities of recombinant soluble TMPRSS2 was assessed using BoC-QAR-AMC fluorogenic substrate over time at 33°C and 37°C. Representative measurements at 33°C (left) and 37°C (middle) are shown. Activity at 33°C and 37°C of 6 ng of recombinant TMPRSS2 is plotted (right). Data are mean ±SD of 4 independent experiments. **c Spike surface expression is unchanged at 33°C and 37°C.** 293T cells were transfected with HKU1 A/B or SARS-CoV-2 Wuhan Spike expression plasmids and incubated at 33°C or 37°C for 24 h. Cells were stained with a pan-coronavirus anti-Spike antibody (mAb10) and analyzed by flow cytometry. Data are mean ± SD of 3 independent experiments. Statistical analysis: Unpaired t-test. **d Spike driven cell-cell fusion is reduced at 33°C.** 293T cells expressing either GFP 1-10 were transfected with HKU1 A/B or SARS-CoV-2 Wuhan Spike. GFP11 cells were transfected with TMPRSS2 or ACE2 and incubated at 37°C. 24 h post transfection, cells were mixed at a 1:1 ratio and incubated at 33°C or 37°C for 2 h 30. Fusion was quantified by measuring the GFP area. Representative images (left) and quantifications (right) are shown. Scale bar = 200 μm. Data are mean ± SD of 3 independent experiments. Statistical analysis: Unpaired t-test. *p<0.05, **p<0.01. **e HKU1 A or B virions are unstable at 37°C.** HKU1 A or B virions were incubated at indicated times and temperatures at 5% CO₂ in a humidified atmosphere before infection of Caco-TMP2+ cells. Infected cells were scored after 24 h. The infectivity of virions after different incubation times, normalized to virions incubated at 4°C for 4 h, is plotted. A representative experiment is shown. Data are mean ± SD of 3 technical replicates. **f Infectivity of virions after 4 h of incubation at indicated temperatures.** Data are mean ± SD of 5 independent experiments. One-way ANOVA. ns = non-significant, *p<0.05, **p<0.01.

**Supplemental Figure 11.**
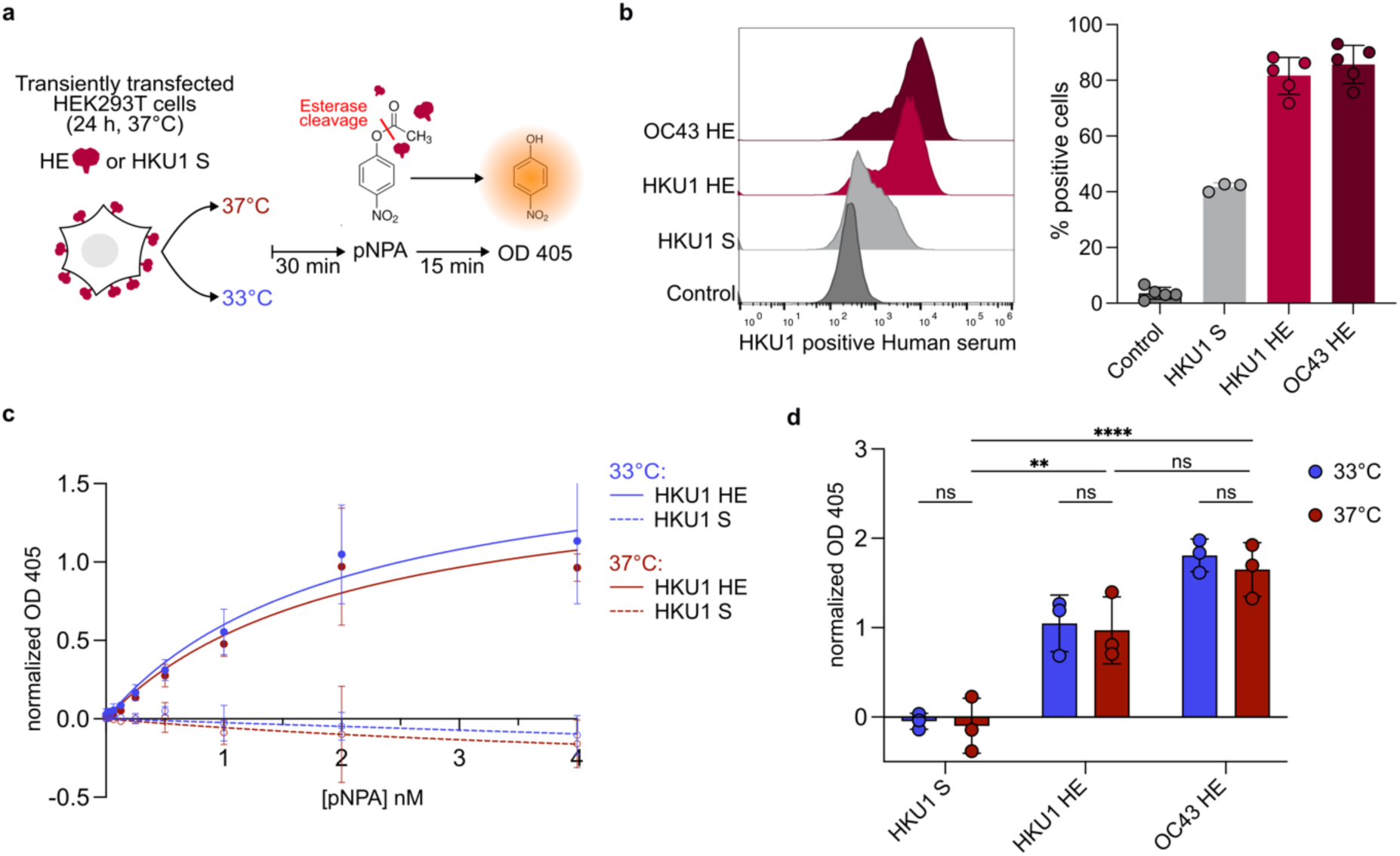
Effect of temperature on HE activity. **a** Protocol for measuring HE esterase activity. Cells were transiently transfected with HKU1 or OC43 HE plasmids or HKU1 S as a negative control. After 24 h, the HE substrate pNPA was added to the cells and OD 405 nm was measured after 15 min incubation at 33° or 37°C. **b** HKU1 and OC43 HE expression is detected by staining with an HKU1 positive Human serum. About 80% of cells are HE positive 24h post-transfection. Data are mean ± SD of 5 independent experiments. **c** The OD 405 nm of HKU1 HE or HKU1 S transfected cells was measured after 15 min incubation with serial dilutions of pNPA, at 33° or 37°C. Normalized OD 405 nm over time is plotted. HKU1 HE exhibits dose-dependent esterase activity toward pNPA. Negative control (HKU1 S) did not exhibit any esterase activity. Data are mean +/- SD of 3 independent experiments. **d** Esterase activity assessed with 2 nM of pNPA. HKU1 and OC43 HE esterase activities are similar at 33 and 37°C. Data are mean ± SD of 3 independent experiments. Statistical analysis: Two-way ANOVA.

**Supplemental Figure 12.**
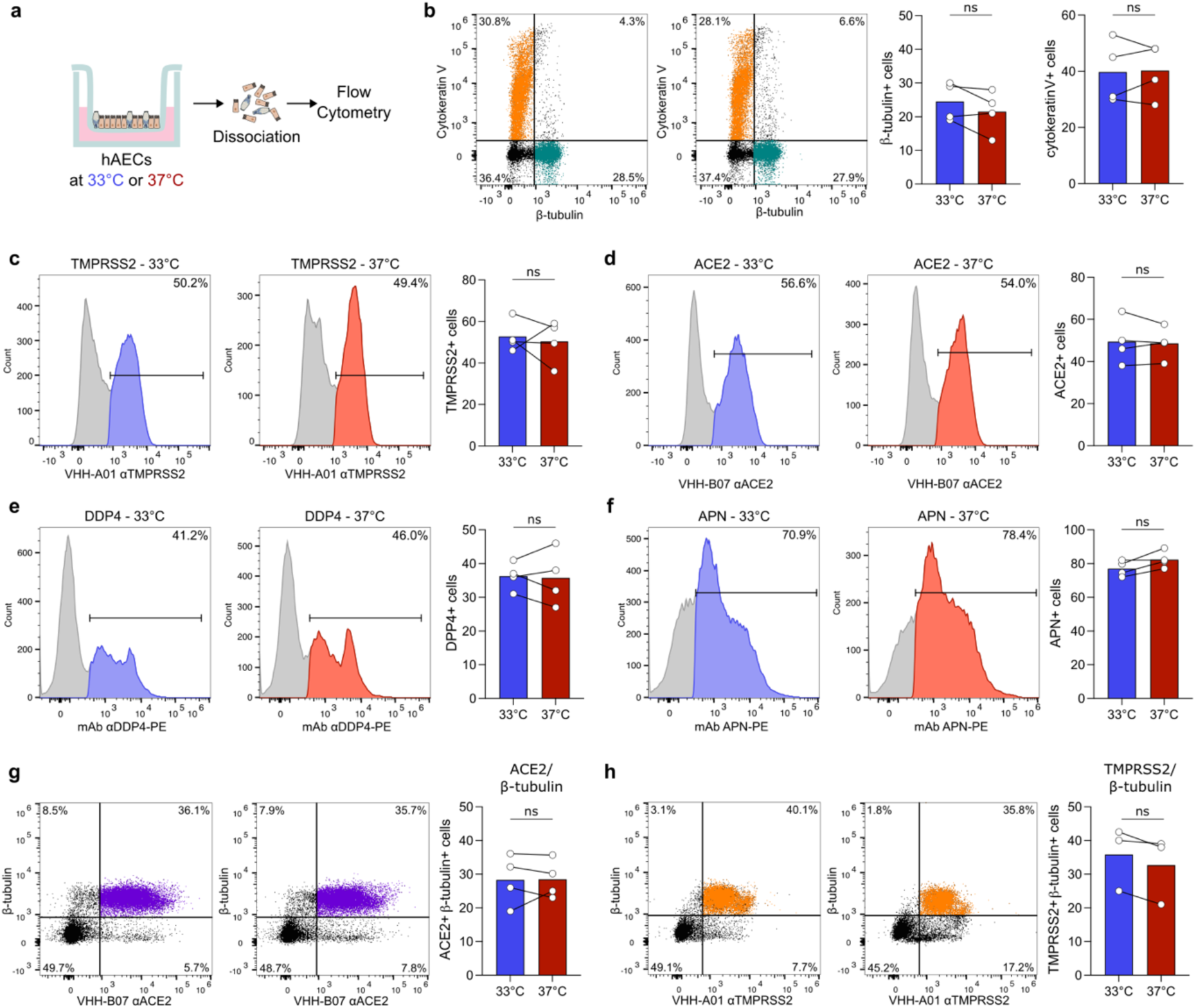
Temperature does not alter coronavirus receptor expression in primary hAECs. **a Protocol overview.** HAECs were maintained at 33°C or 37°C for 72 h, dissociated, stained and analyzed by flow cytometry. **b Percentage of ciliated cells is not changed at 33°C.** Representative dot plots of β-tubulin and cytokeratin V double staining (Left) and quantification of β-tubulin and cytokeratin V positive cells (Right). Data are mean of 4 independent experiments, paired samples are connected by a line. **c-f Temperature does not alter coronavirus receptor surface expression.** Representative dot plots (Left) and quantification of cells positive for TMPRSS2 (C), ACE2 (D), DDP4 (E) and APN (F) (Right). Data are mean of 4 independent experiments, paired samples are connected by a line. **g-h Percentage of ciliated cells expressing ACE2 (g) or TMPRSS2 (h).** Representative dot plots of ACE2/TMPRSS2 and β-tubulin double staining (Left) and quantification of ACE2/TMPRSS2 and β-tubulin double positive cells (Right). Data are mean of 3-4 independent experiments, paired samples are connected by a line. Statistical analysis: Paired t-test. ns = non-significant.

**Supplemental Figure 13.**
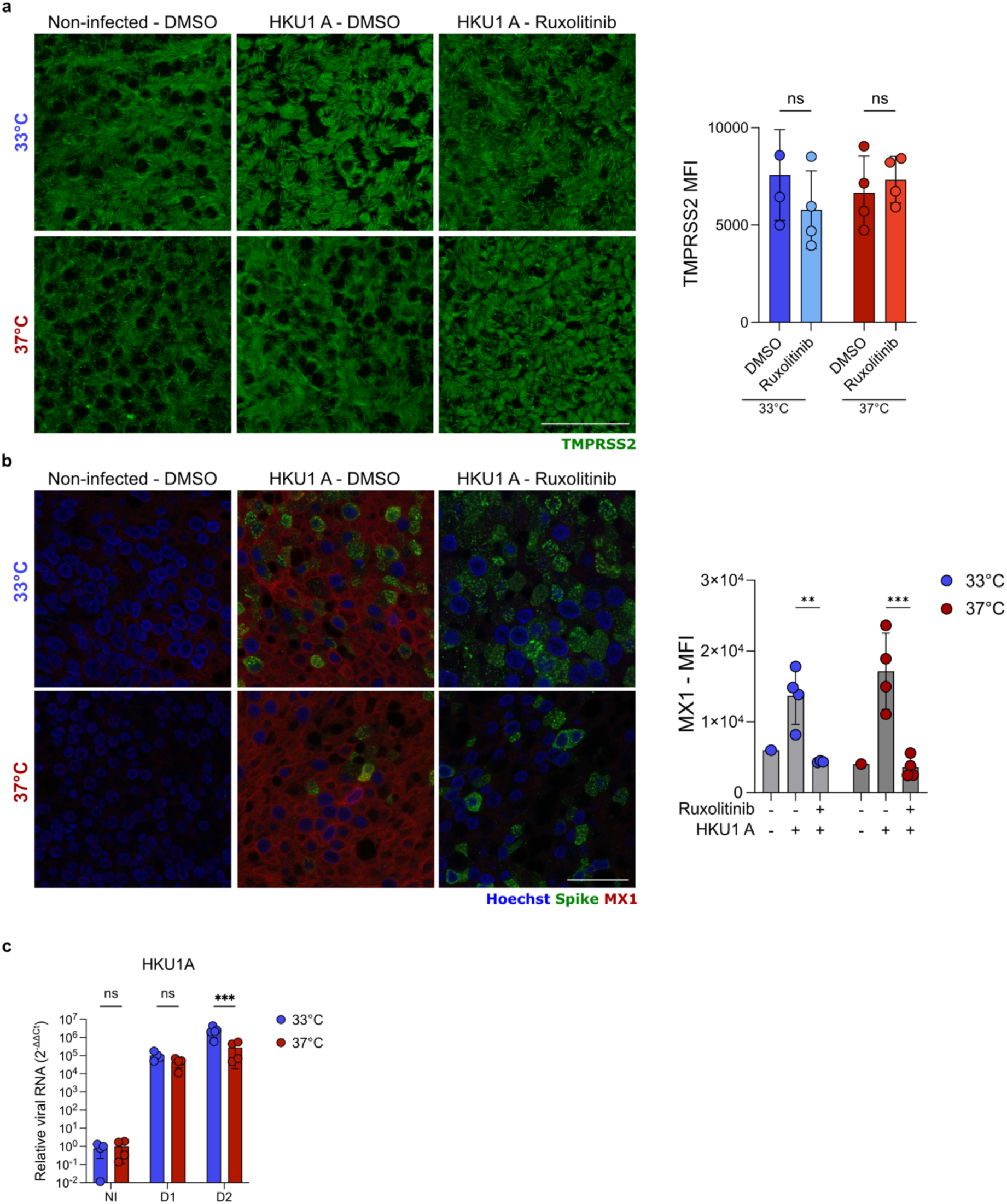
a Temperature and ruxolitinib treatment do not affect TMPRSS2 expression. Representative images (Z-Projection) of TMPRSS2 staining on hAECs at day 4 pi (left). MFI was quantified by ImageJ (right). Each datapoint represents the mean of 4 images. Data are mean ± SD of 4 independent experiments. Statistical analysis: Two-Way ANOVA ns = non-significant. Scale bar = 50 µm. **b MX1 induction upon infection is inhibited by ruxolitinib.** Representative images of MX1 staining on hAECs at day 4 pi (left). Mean fluorescence intensity (MFI) was quantified by ImageJ (right). Each datapoint represents the mean of 4 images. Data are mean ± SD of 4 independent experiments. Statistical analysis: Two-Way ANOVA ** p<0.01, ***p<0.001. Scale bar = 50 µm. **c Intracellular viral RNA quantification of Figure 3i**. Relative quantity of HKU1 RNA normalized to GAPDH endogenous control and non-infected control is plotted. Data are mean ± SD of 4 independent experiments. Statistical analysis: Two-Way ANOVA. ns= non-significant, ***p<0.001.

**Supplemental Figure 14.**
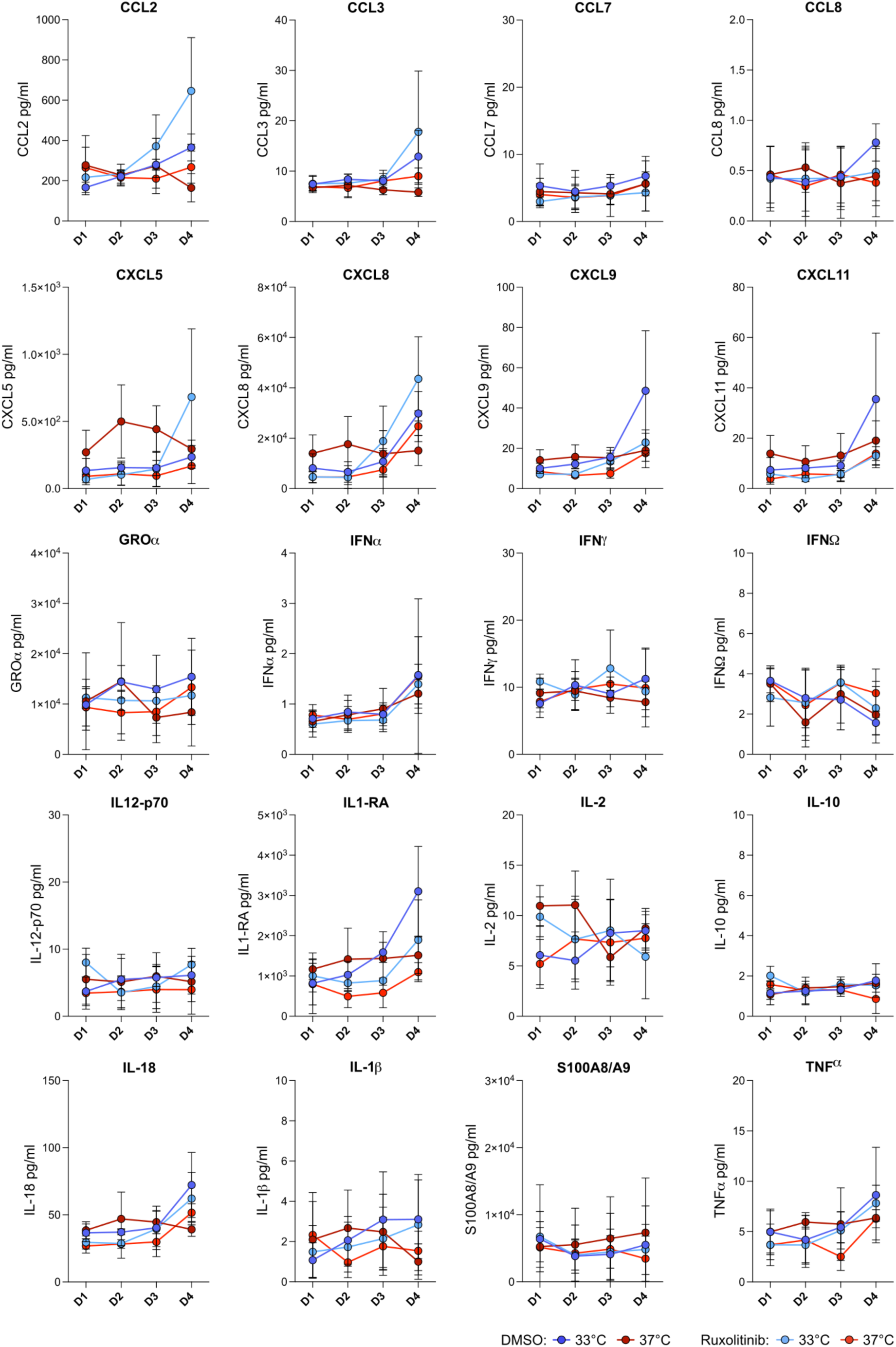
Additional luminex multiplex assay results. Cytokines in the apical compartment of hAECs were quantified over time by Luminex multiplex assay at day 1, 2, 3 and 4. The cytokines are indicated above each graphic. Data are mean ± SD of 4 independent experiments.

**Supplemental Figure 15.**
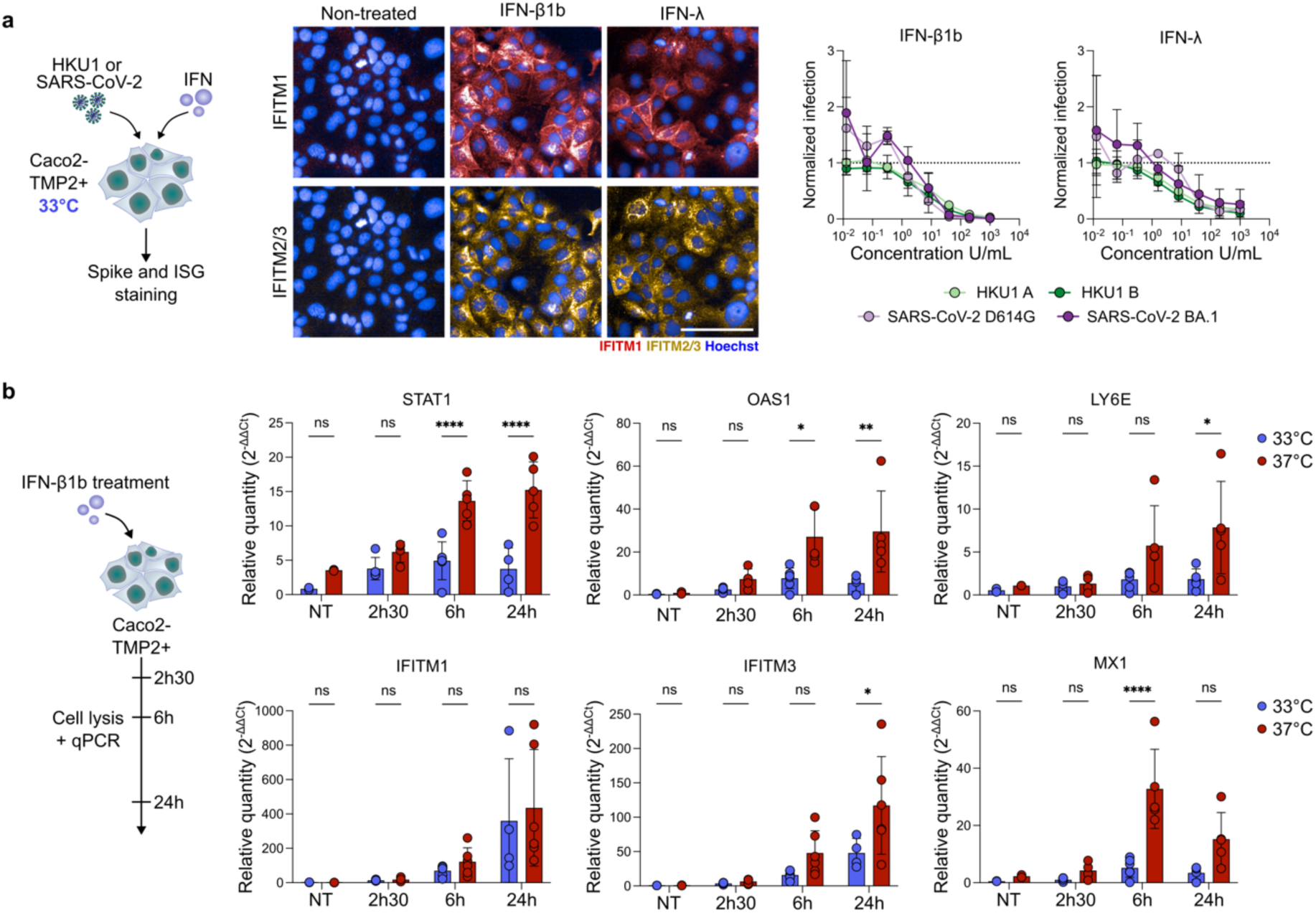
Effect of temperature on ISG induction in Caco2-TMP2+ cells. **a** Caco2-TMP2+ cells were pre-treated with a range of IFN-β1b or IFN-λ for 24 h at 37°C and infected with HKU1 A or B and SARS-CoV-2 (D614G or BA.1 variants). The IFN-β1b or IFN-λ 100 U/ml condition was stained for IFITM1 and IFITM2/3 and imaged. Representative images are shown (middle). IFITM1 (Red), IFITM2/3 (Yellow) and Hoechst (Blue). Scale bar = 100 µm. Spike area was quantified and normalized to non-treated control (right). Data are mean ± SD of 4 (HKU1 A/B) or 2 (SARS-CoV-2) independent experiments. **b Delayed ISG response at 33°C in Caco2-TMP2+ cells.** Caco2-TMP2+ cells were treated with IFN at 100 U/ml and incubated at 33°C or 37°C. Cells were lysed at 2 h 30, 6 h and 24 h post treatment, cellular RNA was extracted and analyzed by RT-qPCR. Relative quantity (2^-ΔΔCt^) normalized to GAPDH endogenous control and non-treated condition is plotted. Data are mean ± SD of 4 independent experiments. Statistical analysis: Two-Way ANOVA * p<0.05, **p<0.01, ****p<0.0001.

**Supplemental Figure 16.**
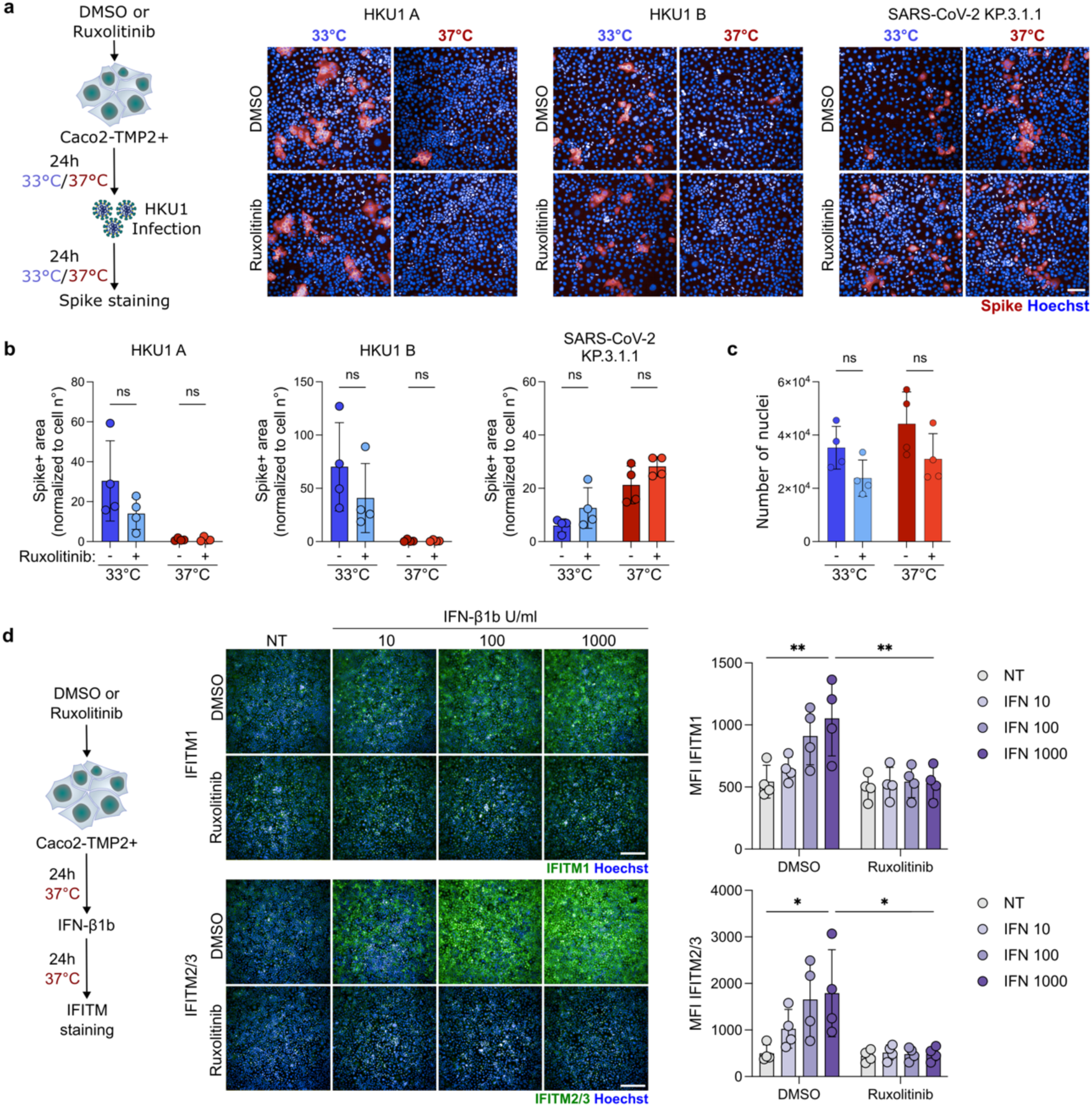
Ruxolitinib does not rescue HKU1 replication at 37°C in Caco2-TMP2+ cells. **a** Caco2-TMP2+ cells were incubated with ruxolitinib (10 µM) or DMSO as a control, for 24 h at 33°C or 37°C before being infected with HKU1 A or B or SARS-CoV-2 (KP.3.1.1 variant). Representative images of Caco2-TMP2+ cells infected with HKU1 A, B or KP.3.1.1 and stained for Spike (red) and Hoechst (Blue). Scale bar = 100 µm. **b** Image quantification of Spike area, normalized to the number of nuclei. **c** Quantification of nuclei in non-infected treated conditions. (B-C) Data are mean ± SD of 4 independent experiments. Statistical analysis: One-Way ANOVA, ns = non-significant. **d** Caco2-TMP2+ cells were incubated with ruxolitinib (10 µM) or DMSO as a control, for 24 h at 37°C before being treated with indicated doses of IFN-β1b, stained for IFITM1 or IFITM2/3 and imaged. Representative images of IFITM1-3 staining in Caco2-TMP2+ cells (middle). IFITM1-3 (Green), Hoechst (Blue). Scale bar = 300 µm. Quantification of IFITM1 or IFITM 2/3 MFI in treated Caco2-TMP2+ (right). Data are mean ± SD of 4 independent experiments. Statistical analysis: Two-Way ANOVA, * p<0.05, **p<0.01.

**Supplemental Figure 17.**
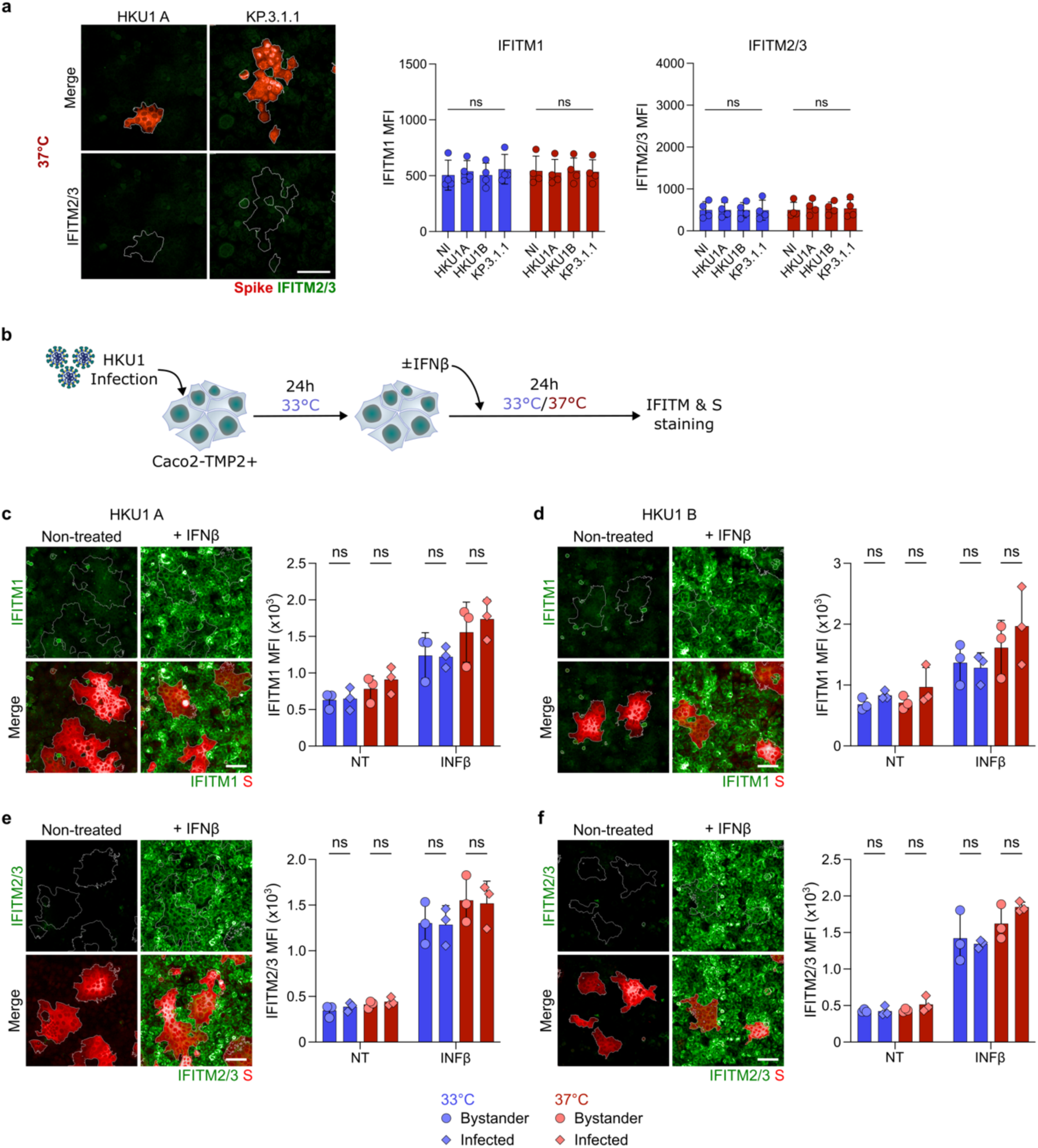
**a** Representative images of IFITM2/3 staining in HKU1 A or KP.3.1.1 infected Caco2-TMP2+ cells (left). S (red), IFITM2/3 (Green). White outline represents Spike positive area. Scale bar = 100 µm. Quantification of IFITM1 or IFITM 2/3 MFI in non-infected and infected Caco2-TMP2+ cells at 33°C (Blue) or 37°C (Red). Data are mean ± SD of 4 independent experiments. Statistical analysis: One-Way ANOVA, ns = non-significant. **b HKU1 infection does not prevent IFN-β1b induced IFITM induction.** Caco-TMP2+ were infected with HKU1 A or B at 33°C for 24h and then treated with IFN-β1b (1000 U/ml) and placed at 33°C or 37°C for an additional 24h. S, IFITM1 and IFITM2/3 were stained and plates were imaged. **b-e** Quantification of IFITM1 (**c-d**) or IFITM2/3 (**e-f**) mean fluorescence intensity in HKU1 A (**c, e**) or B (**d, f**) infected and bystander cells. Left: Representative images. Right: Mean fluorescence intensity quantification of IFITMs. Data are mean ± SD of 3 independent experiments. Statistical analysis: Two-way ANOVA.

**Supplemental Figure 18.**
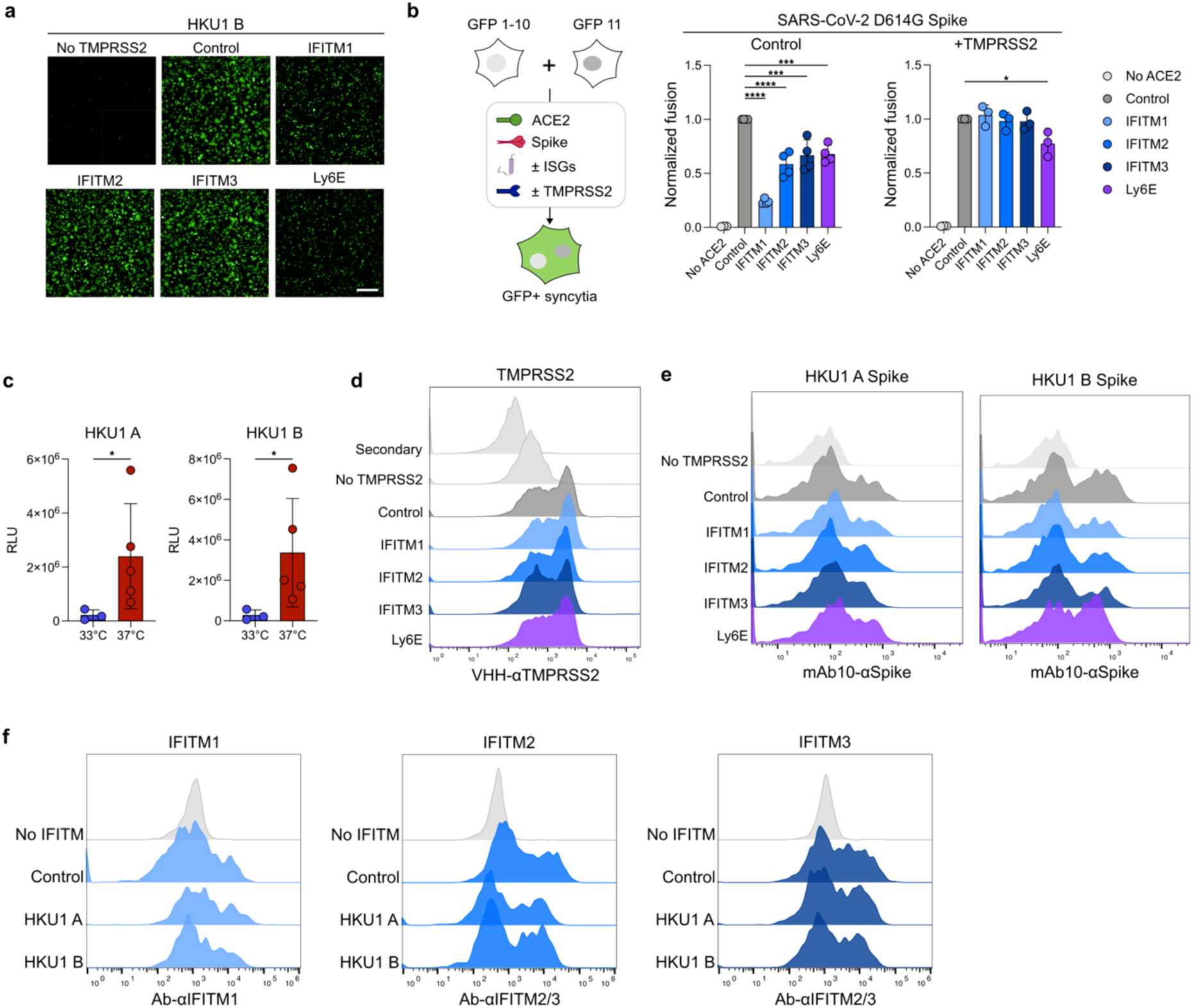
a Representative images of HKU1 B fusion (Figure 4B). b Impact of IFITMs, LY6E and TMPRSS2 on SARS-CoV-2 Spike mediated cell-cell fusion. 293T-GFP1-10 and -GFP11 cells (1:1 ratio) were co-transfected with SARS-CoV-2 D614G Spike, along with TMPRSS2, IFITM, LY6E or control plasmids. Representative images are shown (middle). Scale bar: 500 µm. Cell fusion was quantified by measuring the GFP+ area by high-content imaging after 18 h (right). Data are mean ± SD of 4 independent experiments. Statistical analysis: One Way ANOVA with Tukey’s multiple comparisons * p<0.05, **p<0.01, ***p<0.001, ****p<0.0001. **c Luciferase signal generated by lentiviral pseudovirus is reduced at 33°C.** Non-normalized data of non-treated conditions of Figure 4A showing a reduced RLU signal at 33°C. Data are mean ± SD of 5 (37°C) or 3 (33°C) independent experiments. Statistical analysis: * p<0.05. **d Surface levels of TMPRSS2 in cells expressing ISGs and used for lentiviral pseudovirus infection (Figure 4a).** Cells were stained for TMPRSS2 using VHH-A01-Fc and analyzed by flow cytometry. Representative histograms are shown. **e Surface levels of Spike in cells expressing ISGs and used for cell-cell fusion experiment (Figure 4b).** Cells were stained for HKU1 Spike using mAb10 and analyzed by flow cytometry. Representative histograms are shown. **f Surface levels of IFITMs in cells expressing HKU1 Spikes and used for cell-cell fusion experiments (Figure 4b).** Cells were stained for IFITM1 or IFITM2/3 and analyzed by flow cytometry. Representative histograms are shown.

## Notes

### Competing Interest Statement

The authors have declared no competing interest.

